# Comprehensive Analysis of SARS-CoV-2 Spike Evolution: Epitope Classification and Immune Escape Prediction

**DOI:** 10.1101/2024.12.06.627164

**Authors:** Natália Fagundes Borges Teruel, Matthew Crown, Ricardo Rajsbaum, Matthew Bashton, Rafael Najmanovich

**Affiliations:** Department of Pharmacology and Physiology, Faculty of Medicine, Université de Montréal, Montreal, Canada; Hub for Biotechnology in the Built Environment, Department of Applied Sciences, Faculty of Health and Life Sciences, Northumbria University, Newcastle upon Tyne, UK; Center for Virus-Host-Innate Immunity and Department of Medicine; Rutgers Biomedical and Health Sciences, Institute for Infectious and Inflammatory Diseases, Rutgers University, Newark, New Jersey, United States of America

**Keywords:** SARS-CoV-2, Spike protein, structural biology, epitopes, immune recognition, conformational dynamics

## Abstract

The evolution of SARS-CoV-2, the virus responsible for the COVID-19 pandemic, has produced unprecedented numbers of structures of the Spike protein. This study presents a comprehensive analysis of 1,560 published Spike protein structures, capturing most variants that emerged throughout the pandemic and covering diverse heteromerization and interacting complexes. We employ an interaction-energy informed geometric clustering to identify 14 epitopes characterized by their conformational specificity, shared interface with ACE2 binding, and glycosylation patterns. Our per-residue interaction evaluations accurately predict each residue’s role in antibody recognition and as well as experimental measurements of immune escape, showing strong correlations with DMS data, thus making it possible to predict the behaviour of future variants. We integrate the structural analysis with a longitudinal analysis of nearly 3 million viral sequences. This broad-ranging structural and longitudinal analysis provides insight into the effect of specific mutations on the energetics of interactions and dynamics of the SARS-CoV-2 Spike protein during the course of the pandemic. Specifically, with the emergence of widespread immunity, we observe an enthalpic trade-off in which mutations in the receptor binding motif (RBM) that promote immune escape also weaken the interaction with ACE2. Additionally, we also observe a second mechanism, that we call entropic trade-off, in which mutations outside of the RBM contribute to decrease the occupancy of the open state of SARS-CoV-2 Spike, thus also contributing to immune escape at the expense of ACE2 binding but without changes on the ACE2 binding interface. This work not only highlights the role of mutations across SARS-CoV-2 Spike variants but also reveals the complex interplay of evolutionary forces shaping the evolution of the SARS-CoV-2 Spike protein over the course of the pandemic.

## 1 Introduction

The severe acute respiratory syndrome coronavirus 2 (SARS-CoV-2) virus, responsible for the global COVID-19 pandemic,^1–3^ has been the subject of extensive scientific scrutiny, particularly regarding its Spike protein, which plays a critical role in viral entry into host cells. The ability of the SARS-CoV-2 Spike protein (referred simply as Spike from here on) to bind to the Angiotensin-converting enzyme 2 (ACE2) receptor,^4, 5^ its immune recognition,^6^ and its glycan shield^7^ have made it a focal point of research, leading to an unprecedented amount of structural data.^8^ This wealth of information has enabled researchers to study the molecular mechanisms governing viral infectivity and immune evasion, using both experimental and computational approaches.

While computational and experimental studies have provided detailed insights into specific aspects of the Spike protein related to binding^9, 10^ and dynamics,^11–14^ most studies often do so in isolation, focusing on individual aspects rather than the broader functional context. Conversely, epidemiological studies tend to capture the overall effects of these mechanisms on viral spread, yet lack the granularity needed to disentangle the contributions of multiple mechanisms to the observed outcomes.

The central role of the Spike protein in mediating viral entry also makes it a prime target for immune recognition and therapeutic intervention.^6^ Its surface exposure and dynamic conformational changes create opportunities for immune recognition but also pose challenges to the immune system due to the presence of protective mechanisms like glycan shielding and conformational masking. Key regions, such as the receptor-binding domain (RBD) and the RBM, are critical for neutralizing antibody responses and are often subject to evolutionary pressures that balance receptor binding efficiency with immune evasion.^15^ Dissecting how these regions evolve under such pressures provides valuable insights into the interplay between viral infectivity and immune escape.

A crucial step in understanding this interplay involves the characterization of epitopes. Beyond their role in antibody recognition, epitopes are defined by their accessibility, which is influenced by glycan shielding and conformational flexibility, as well as by the shared binding interface with ACE2 and the constraints to the mutational landscape that this necessary interaction imposes. Numerous studies have employed different experimental and computational methodologies to identify epitopes,^16–27^ with a predominant focus on the RBD.^16, 18–22, 26, 27^ However, while experimental methodologies are often constrained by their high cost, time-consuming nature, and complexity — making frequent updates challenging — many studies based on structural analyses were limited by the availability of structural data at the time and the absence of reliable high-throughput methods for evaluations of energetics of molecular interactions.

Computational approaches are essential for understanding viral evolution, both in the unification and evaluation of viral genetic data^28–31^ and, in the context of structural analysis, in characterizing specific mechanisms and exploring potential mutations and their effects.^9, 11, 13, 32^ Such computational approaches often employ modeling techniques to introduce single or multiple mutations, investigate their functional effects, and explore the dynamic range of motion and accessibility across different conformational states.^10, 13^ However, these methods inherently introduce biases due to limitations in the modeling process,^33, 34^ affecting subsequent analyses.^35, 36^ Studies based on experimental structures — that are also subject to biases^37^ — often provide a limited view of functional aspects, as they typically rely on a single or few structures^38–40^ that fail to capture the full variability and dynamic nature of a moving protein, a limitation that can be seen when comparing large numbers of observations. Meanwhile, a vast amount of experimentally solved structural data remains underutilized, primarily due to limitations of computationally expensive approaches in evaluating extensive datasets and conducting detailed per-residue analyses.^10^ By employing methodologies suitable to high-throughput applications, we can leverage this extensive structural data for more comprehensive and accurate assessments, averaging out experimental biases inherent in protein structure determination.

In this study, we perform a comprehensive structural analysis of 1,560 Spike protein structures with respect to interactions with antibodies and the receptor ACE2, conformational dynamics and glycan coating. We aim to provide a broad understanding of how various structural features contribute to the function and evolution of the Spike protein. To do so, we introduce a data-driven epitope classification system and a method for mapping other complexes into one of the 14 epitopes identified wihtin this study. Uniquely, our methodology allows us to approximate the enthalpy of pairwise-interacting complexes as the sum of pairwise residue-level interactions in a high-throughput fashion and predict the experimentally determined immune escape potential of individual mutations. This approach is essential to understand the complex interplay between different aspects of the Spike protein, offering new perspectives on its role in viral evolution.

## 2 Methods

### 2.1 Structure selection and modeling

We utilized all Spike protein structures available in the RCSB Protein Data Bank until October 2nd, 2023. These structures were annotated based on their complexes — either with antibodies, with the receptor ACE2, or unbound — as well as based on the presence of glycans, the variant they represent, and, in the case of full homotrimeric structures, the conformational state of each Spike monomer. We utilized pairs of structures that had the exact same sequence, were published together, and represented the closed state and the one-RBD-open state to estimate open state occupancy as described in Teruel *et al.*^13^ For these pairs of structures, we rebuilt the missing loops using Modeller.^41^ The open state estimation of occupancy is calculated only for variants for which the required structures above are present in the PDB.

### 2.2 Evaluation of interaction energies

We used the software Surfaces^35^ to evaluate the interaction free energy of complexes of Spike and antibodies, Spike and ACE2, or the interactions of glycans present in the structures. Surfaces was previously shown to predict the effects of Spike mutations on the binding affinity to the receptor ACE2 as accurately as current state-of-the-art methods based on molecular dynamics simulations at a fraction of the required computational time.^35^ Surfaces generates a matrix of pairwise pseudo-energetic contributions to the enthalpy of binding. We collapsed the matrix of pairwise interactions into a vector representing the sum of all interaction enthalpies for every Spike residue. In the case of interactions with more than one chain of Spike, we focus on the interacting chain that contributes the most to the total enthalpy. We then aligned the vectors to a reference sequence in order to correct incorrectly numbered residues or adjust the numbering in cases of structures with insertions and deletions. For both per-residue evaluations and full interface interactions, we refer to the pseudo-energy calculated by Surfaces as binding affinity, as it is scaled to represent the free energy of binding in units of kcal/mol.

### 2.3 Antibody clustering

Given a protein-protein interaction interface of a Spike-Antibody complex, we define a reference point as the geometric average position of all interacting residues, weighted by their contribution to the overall interface enthalpy as calculated by Surfaces:

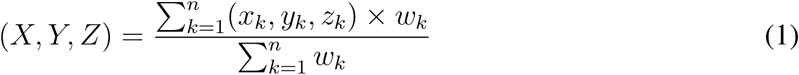

where (*X, Y, Z*) represent the coordinates of the reference point, (*x_i_, y_i_, z_i_*) are the coordinates of each residue within the interface, with its corresponding contribution to the enthalpy *w*; *n* represents the number of interface residues.

Hierarchical clustering based on the pairwise Euclidean distance of these average points was used to cluster the 2,032 antibody interactions, from 942 structures of Spike-antibody complexes (Fig S1). This is, to our knowledge, the first instance of using interaction energies to bias geometric clustering, adjusting the relevance of the interaction interface based on the importance of the per-residue contacts. This approach resulted in 14 clusters, represented by their respective weighted average points, that are biased towards the positions with the strongest interactions in each epitope. We applied principal component analysis (PCA) to visualize the clustering results. This enabled us to project the points into a 2D space based on their weighted average interactions, simplifying the interpretation of the clusters.

### 2.4 Vector sorting

We calculate the distance of the average points of 2,032 antibody interactions, as described by equation 1, to each of the 14 reference points representing the characterized epitopes. This method provides greater clarity than considering all pairwise distances, especially in differentiating clusters that are very close in space, such as epitopes 4 and 5.

Employing the same strategy described by equation 1, the 209 ACE2-Spike interaction interfaces, based on 146 complex structures, are mapped to unique epitopes by finding the epitope reference point closest to the average point of interaction for the particular ACE2-Spike interface. The equation above is also used to analyze single residues, particularly residues in positions that undergo conformational changes or glycosylated residues. In the case of single residues, its reference point as calculated by the equation above reduces to the coordinates of the given residue atoms — and likewise for entire interfaces, this reference point is mapped to the epitope closest in space.

### 2.5 Occupancy evaluation

We employed the NRGTEN as described by Mailhot *et al.*^42^ to evaluate the occupancy of conformational states of the Spike protein. This method builds upon previous work on the structural dynamics of the Spike protein,^13^ incorporating Markov Chain modeling to calculate conformational state occupancies.

In the Markov model, each conformation of the Spike protein is represented as a state, with transition probabilities between states calculated using the ENCoM model. A constant *k* is added to all states, representing the probability of remaining in that state. To ensure that the transition probabilities from each state sum to 1, these values are normalized after the addition of *k*. This approach results in a unique equilibrium solution, which provides the occupancy values for the conformational states.

For this study, we performed a new parametrization of the model using data from six structural pairs representing different variants of the Spike protein: *wild-type* (PDB 7KDG and 7KDH^43^), D614G (PDB 7KDI and 7KDJ^43^), Beta (PDB 7WEV and 7VX1^44^), Kappa (PDB 7VXI and 7VXE^44^), Delta (PDB 7W94 and 7W92^45^), and Omicron (PDB 7WK2 and 7WVN^40^). The occupancy values associated to these structures were used to optimize the parameters *k* and γ for our system. The selected constants, *k* = 0.1 and γ = 0.1, yielded a Pearson’s correlation of 0.657 with the experimental occupancy values, showing good predictive accuracy.

Using this re-parameterized model, we derived a linear transformation to relate the calculated occupancy difference between open and closed states to the experimental values:

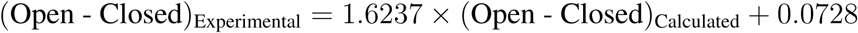

This transformation allows us to quantitatively compare the model’s predictions with observed experimental data with a robust mechanism for evaluating the conformational dynamics of the Spike protein. This method was employed to 180 homotrimeric structures of Spike, organized in pairs, as described above (see Structure selection and modeling).

### 2.6 Longitudinal Analysis of Antibody Escape and Vibrational Difference Score

Global longitudinal genomic data analysis of SARS-CoV-2 was performed using SPEAR v2.1^46^ against all consensus genome sequences deposited in GenBank^47^ (accessed 2024-08-17) and marked as complete. A total of 3,048,694 samples were available for analysis with SPEAR. SPEAR integrates deep mutational scanning data on antibody recognition and predicted Vibrational Difference Score (VDS) into a number of scores which can be tracked longitudinally.

From the GenBank dataset (n=3,048,694), 2,994,442 met QC cutoffs and were analyzed using SPEAR. Some samples do not have a valid collection date (for example Omicron samples occurring in 2020) or samples were specific to a particular year (which is not informative for longitudinal analysis). Samples were filtered to only include lineages where a collection date was present and was less than 100 days prior to lineage designation, and had a sample date specific to a year and month at a minimum, resulting in a total of 2,841,216 samples being retained, spanning from December 2019 to August 2024. Samples were assigned to 3,353 different Pangolin lineages, corresponding to 43 Nextstrain clades and 13 Variants of Concern (VOCs) (and a Not VOC class). From all 2,994,442 analyzed samples, representative VCF files for 2,941 lineages were created and analyzed (requiring a minimum of 5 samples assigned to a lineage for analysis reduces the total number of lineages), using the SPEAR `utilities-representative` function. For a mutation to be included as representative of a lineage, it must occur within 50% of all samples assigned to a lineage. The lineage representative VCF files were then re-analyzed using SPEAR in `VCF` mode in order to explore the functional effects characteristic mutations of these lineages exhibit.

### 2.7 Determination of factors influencing Trade-off

The trade-off analyses compare occupancy, antibody binding, and receptor binding calculations pairwise for particular mutations or sets thereof, such as variants. We utilized bootstrapping in order to account for differences in the amount of available data. For each pair, we built four datasets, containing data points from structures representing *wild-type* and each variant for the two functional properties. From these datasets, we generated 500 bootstrap samples and calculated the difference between the variant and *wild-type* for the mean value of each functional property.

## 3 Results and Discussion

### 3.1 Antibody recognition

Epitope characterization plays a key role in understanding immune recognition of the SARS-CoV-2 Spike protein and the mutations that disrupt antibody interactions. Accurately defining epitopes is crucial for determining the antibodies that need to be included to assess the impact of mutations on immune escape. Based on 2,032 interaction vectors between Spike and antibodies (See Antibody Clustering), we assigned each complex to one of the 14 different epitopes (Fig 1A,B,D) using their weighted representative point (See Vector sorting).

**Fig 1.**
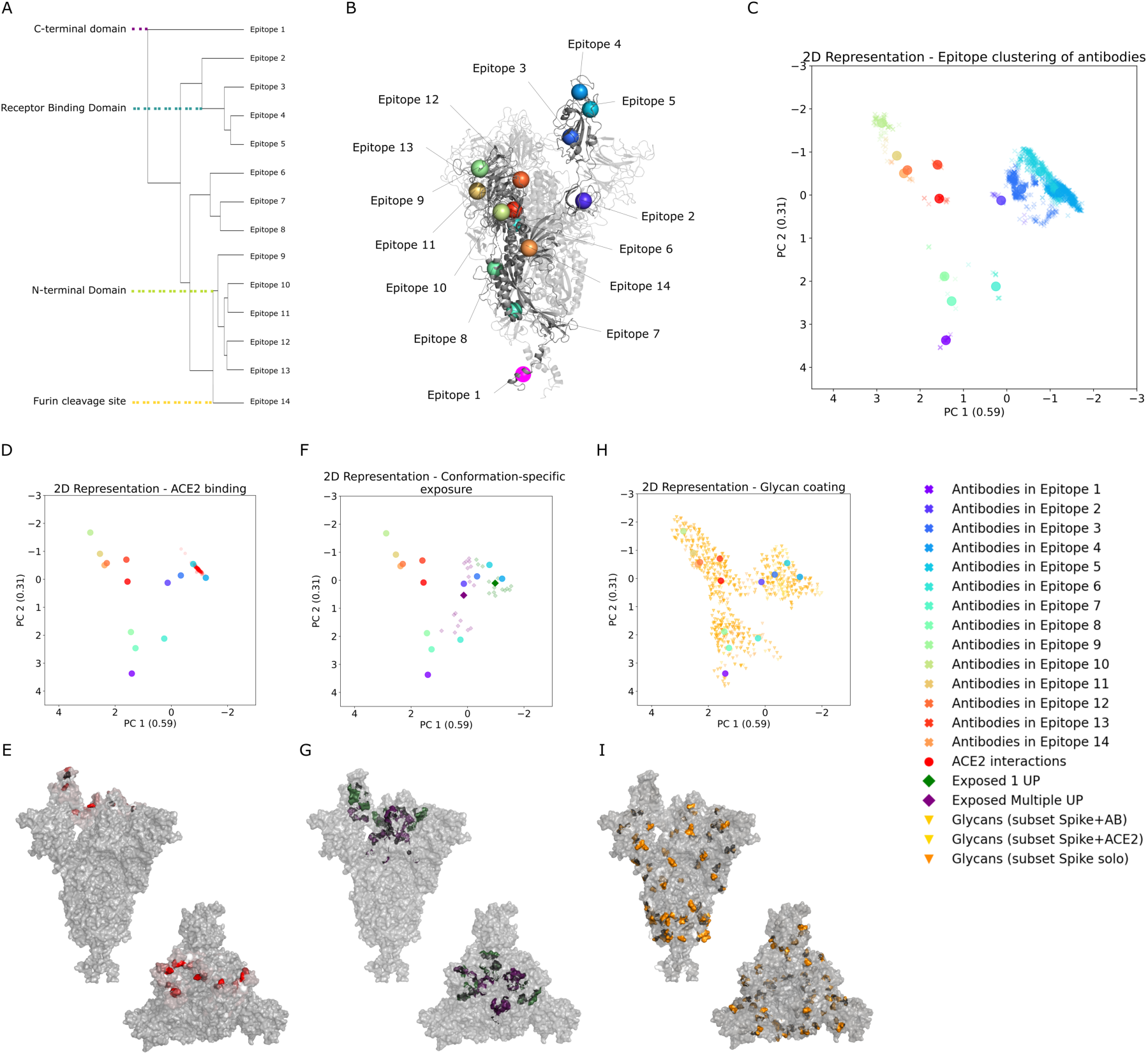
Schematic representation of epitope clustering and structure mapping. (A) Dendrogram representing the hierarchical clustering of SARS-CoV-2 Spike protein epitopes based on their average positions, (B) highlighted as colored spheres, in the open chain of the Spike structure, with each color corresponding to a distinct epitope cluster. (C) Two-dimensional representation of antibody binding data using PCA, with respective variances indicated, showing the clustering of antibodies based on the distance of each average point of interaction to each epitope average position on the Spike protein. (D) The same technique for two-dimensional representation was employed to plot 209 ACE2 binding interactions (red spheres), showing the proximity to each epitope center. (E) We can also see the mapping in the Spike surface of the sum of ACE2 vectors of interaction. (F) Another PCA plot shows residues with conformation-specific exposure, dependent on at least one RBD up (green diamond) or multiple RBD up (purple diamond); the average position of these residues is represented by larger markers for the same two groups. (G) Residues with conformation-specific exposure shown in the Spike surface. (H) PCA representation of glycan interactions for the subsets of structures of Spike in complex with antibodies (gold triangle), Spike in complex with ACE2 (yellow triangle), or unbound Spike (orange triangle). (I) Glycan coating for the unbound Spike subset mapped to the Spike protein surface.

The most common epitopes, 3, 4 and 5 (Fig S2A), describe interactions with the RBD, but with different orientations: Epitope 3 is characterized by strong interactions with positions 377 and 378 (Fig S2E). Epitope 4 is governed by residue 486 (Fig S2F), with interactions encompassing the region of the RBM. The strongest interacting residue in epitope 5 is 346, a residue that is considerably accessible even in closed conformations (Fig S2G).

Epitopes 9, 10, 11, 12 and 13 are localized within the N-terminal Domain (NTD). This domain, located on the external side of the homotrimeric Spike structure, adopts a *µ*-sandwich fold and a forms stable core. One of its sides faces the RBD of the same monomer, while the other side and the top of the domain are close to the RBD of the neighboring monomer. Epitope 9 shows interactions in the external tip of the NTD, namely residue 147 (Fig S2K). Lower to the tip is a part of the NTD that constitutes the epitope 10, with governing residues 214 and 218 (Fig S2L). Epitope 11 characterizes a lateral portion of the NTD, closer to the neighboring chain, mainly residues 173 and 176 (Fig S2M). The other lateral portion of the NTD forms a binding pocket along with the post-RBD area of the same Spike chain, creating epitope 12, with governing residues 85 and 237 (Fig S2N). Epitope 13 is located on the extreme lower part of NTD, and is often characterized by buried peptides, mainly positions 277, 269 and 270 (Fig S2O).

Epitope 1 characterizes the interactions in the C-terminal Domain, mainly residues 1148 and 1156 (Fig S2C). Epitope 2 is characterized by interactions in the external surface of Spike chains, after the RBD, with its main interacting residue being number 537 (Fig S2D). Epitopes 6 (Fig S2H), 7 (Fig S2I), 8 (Fig S2J), and 14 come mainly from structures of antibodies interacting with Spike peptides that are buried into the structure in the context of the entire homotrimeric protein. Another characteristic of epitope 14 is that it describes interactions with the furin cleavage site peptide (Fig S2P). For epitope 8, we also see interactions to the homotrimeric structure in the cases of glycan-mediated antibody binding (Fig S6C). The main interacting positions of these epitopes are described in Table 1.

**Table 1.** Main Spike residues for each epitope. Residues with a mean interaction of -0.75 kcal/mol or lower, ordered from stronger to weaker mean binding affinity.

| Epitope | Main Spike Residues |
| --- | --- |
| 1 | 1156, 1148, 1155, 1152, 1149, 1151, 1157 |
| 2 | 537 |
| 3 | 378 |
| 4 | 486, 489 |
| 5 | 346 |
| 6 | 1008, 1001, 765, 767, 1007, 1000, 772 |
| 7 | 904, 918, 898, 911, 917, 896, 897, 914, 912, 910 |
| 8 | 823, 815, 819, 822, 820 |
| 9 | 147, 248, 249, 246, 144, 146 |
| 10 | 214, 97, 69 |
| 11 | 173, 176, 175 |
| 12 | 237, 85, 83 |
| 13 | 277, 270, 269, 275, 271, 276, 274, 273 |
| 14 | 685 |

An important caveat is that many antibodies interact with more than one Spike chain simultaneously. For the purpose of assigning and defining epitopes, however, we consider only those with the Spike chain with which most significant interactions occur (based on Surfaces calculations). Of the 2,032 antibody interaction vectors, when filtering out crystallization artifacts, there are 338 additional interactions with neighboring Spike chains (Fig S3A). On average, the calculated absolute value of interaction pseudo-energy with these neighboring chains is 7% of that with the primary interacting Spike chain. When classifying the secondary interactions according to the epitopes they target, we see the majority of the interactions with the neighboring RBD, mainly epitope 3 (Fig S3B), but also with the neighboring NTD (Fig S3C). A notable example of dual interaction is the antibody S2M11, which is classified as interacting with epitope 4 but also binds to the neighboring epitope 3 with up to 69% of the binding affinity observed for the primary interacting chain.

Several epitope classifications exist,^16–27^ underscoring the value of epitope classification in SARS-CoV-2 research. However, the classification of Barnes *et al.*^18, 19^ is the most widely adopted as a framework for studying antibody interactions. Although Barnes’ classification offers a concise overview of RBD epitopes, it (and others cited above) lacks practical tools for classifying novel complexes and excludes epitopes outside the RBD. Since the initial establishment of these classes in 2020, the volume of structural data available has grown significantly, allowing for a more robust and data-driven approach to epitope determination that addresses the limitations of existing classifications, including that of Barnes.

Given the widespread use of this classification, we assessed the equivalence of our own categorized epitopes to Barnes’ four classes by analyzing epitopes targeted by a set of well-documented antibodies with experimentally determined structures in complex with the Spike protein. Using Surfaces to calculate per-residue pseudo-energies, we found a strong correlation between residues crucial for each interaction and the total escape per site calculated for the same antibodies based on deep mutational scanning (DMS) results^48, 49^ (Table 2). By examining these metrics across each class and epitope, we observed alignment between Barnes’ classes 1 and 2 with our epitope 4, class 3 with epitope 5, and class 4 with epitope 3 (Fig 2). Notably, while classes 1 and 2 show substantial overlap in escape data characterizing critical residues for antibody recognition, their distinction arises from a secondary interaction with an adjacent Spike chain — a feature not apparent in escape assays due to the experimental use of expressed RBD rather than the full homotrimeric Spike complex, and a distinction our interaction evaluations are able to capture (Fig 2).

**Table 2.** Epitope classification equivalence.

| Antibody | Barnes' Class | Epitope | PDB ID | Pearson's correlation |
| --- | --- | --- | --- | --- |
| S2E12 | 1 | 4 | 7K45, 7K4N, 7R6X | -0.86 |
| REGN10933/casirivimab | 1 | 4 | 6XDG | -0.81 |
| COV2-2196/AZD8895 | 1 | 4 | 7L7D, 7L7E | -0.80 |
| C102 | 1 | 4 | 7K8M | -0.62 |
| LY-CoV555/bamlanivimab | 2 | 4 | 7L3N | -0.88 |
| S2M11 | 2 | 4 | 8DLW, 7K43, 7SO9, 7LXY | -0.76 |
| C002 | 2 | 4 | 7K8S, 7K8T | -0.75 |
| C144 | 2 | 4 | 7K90 | -0.63 |
| C121 | 2 | 4 | 7K8X, 7K8Y | -0.53 |
| REGN10987/imdevimab | 3 | 5 | 8J26, 8J1Q, 7ZJL | -0.75 |
| COV2-2130/AZD1061 | 3 | 5 | 8SUO, 7L7E | -0.75 |
| C135 | 3 | 5 | 7K8Z | -0.73 |
| LY-CoV1404/bebtelovimab | 3 | 5 | 7MMO | -0.69 |
| C110 | 3 | 5 | 7K8V | -0.63 |
| CR3022 | 4 | 3 | 7A5R, 6W41 | -0.66 |
| S2H97 | 4 | 3 | 7M7W | -0.62 |
| COVA1-16 | 4 | 3 | 7LM8, 7S5Q | -0.58 |
| S304 | 4 | 3 | 7JW0, 7X1M, 7R6X | -0.52 |

**Fig 2.**
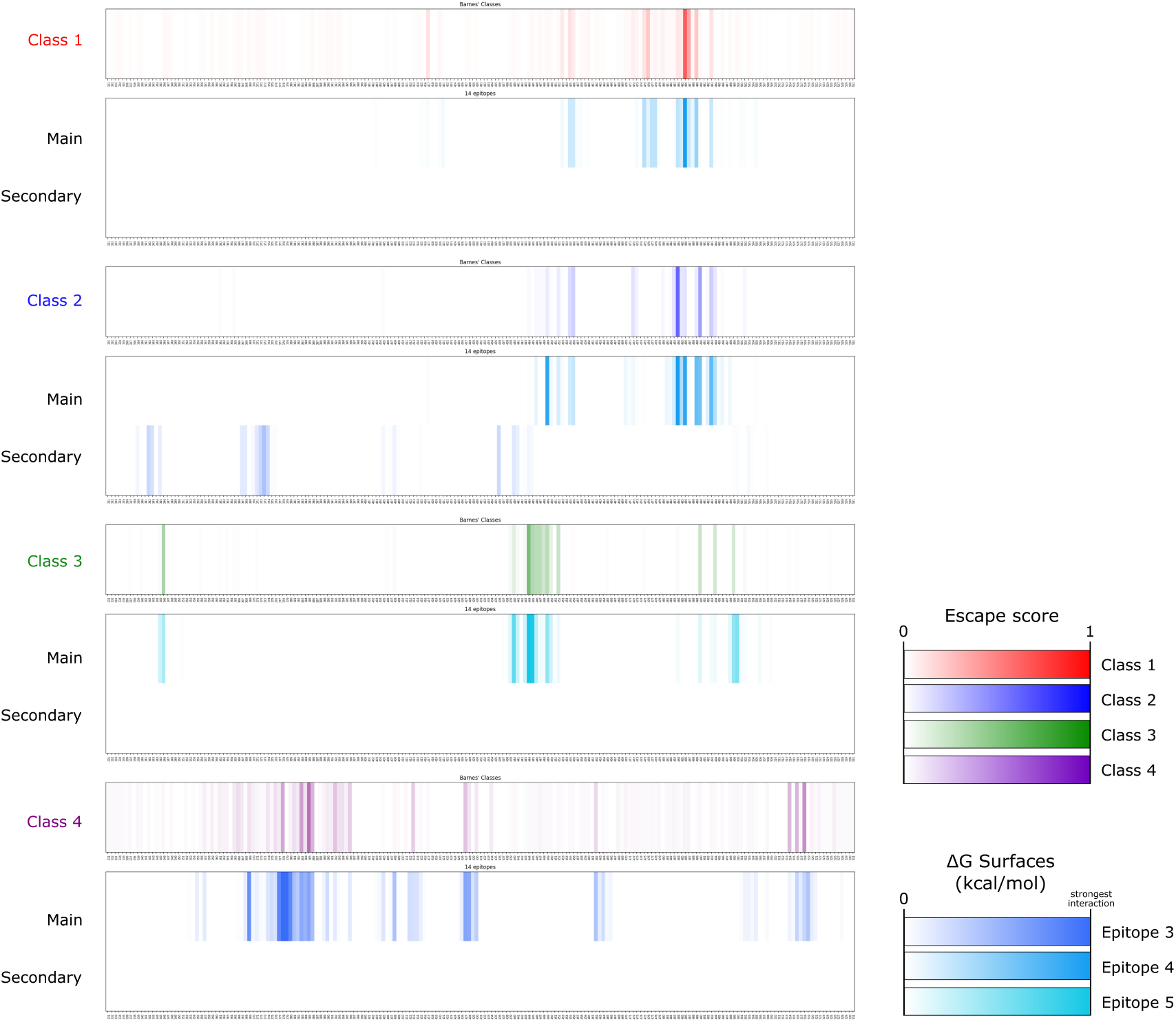
Comparative analysis of epitope classifications. Comparative analysis of epitope classifications between Barnes *et al.* (2020)^18,19^ and our data-driven epitope determination approach. Each row represents an epitope class from Barnes’ classification (Classes 1–4), represented by the site escape scores for each RBD residue, and the respective pseudo-energies calculated via Surfaces, showing interaction strength across different epitopes (Epitopes 3, 4, and 5). Class 1 is represented by escape data from antibodies S2E12, COV2-2196/AZD8895, REGN10933/casirivimab, and C102, and is equivalent to interaction data characterizing main interactions to epitope 4 from structures 7K45, 7K4N, 7R6X, 7L7D, 7L7E, 6XDG and 7K8M. Class 2 is represented by escape data from antibodies LY-CoV555/bamlanivimab, S2M11, C002, C144, and C121, and is equivalent to interaction data characterizing main interactions to epitope 4 and secondary interactions to epitope 3 from structures 7L3N, 8DLW, 7K43, 7SO9, 7LXY, 7K8S, 7K8T, 7K90, 7K8X, and 7K8Y. Class 3 is represented by escape data from antibodies REGN10987/imdevimab, COV2-2130/AZD1061, C135, LY-CoV1404/bebtelovimab, and C110, and is equivalent to interaction data characterizing main interactions to epitope 5 from structures 8J26, 8J1Q, 7ZJL, 8SUO, 7L7E, 7K8Z, 7MMO, and 7K8V. Class 4 is represented by escape data from antibodies CR3022, S2H97, COVA1-16, and S304, and is equivalent to interaction data characterizing main interactions to epitope 3 from structures 7A5R, 6W41, 7M7W, 7LM8, 7S5Q, 7JW0, 7X1M, and 7R6X.

Beyond establishing the equivalence with Barnes’ classes, our evaluation demonstrates that per-residue contributions as calculated with Surfaces can predict DMS-based immune escape calculations. Specifically, experimental immune escape scores^48, 49^ (ranging from 0 to 1) align with per-residue interactions (measured as pseudo-energy, where more negative values indicate stronger interactions). Table 2 shows the correlation between the average per-residue immune escape score and Surfaces’ interaction energy predictions for several instances of different antibodies. The Pearson’s R obtained range from -0.52 to -0.86. It is noteworthy that the average correlations are markedly lower for antibodies in class 3 (Barnes class 4). This alignment emphasizes the predictive accuracy of Surfaces in identifying immune escape potential across different Spike residues and Spike-targeting antibodies.

This comparative evaluation could be extended to all antibodies with available structures in complex with the Spike protein and included in DMS studies, provided that the nomenclature of these antibodies is standardized. As a preliminary assessment, the results we present for the 18 antibodies demonstrate promising predictive accuracy.

The longitudinal plot shown in Figure 3A shows the number of mutations that the 14 characterized epitopes accumulated in the timeline of the pandemic. Starting in the Alpha variant, that did not exhibit significant antibody escape,^50^ we see mutations starting to accumulate on epitope 4, with the substitution N501Y, that presented an increase in overall binding affinity in structural evaluations (Fig 3B). The emergence of Delta coincides with an increase in immune escape, driven by a number of mutations to Spike^51^ including L452R,^52^ part of epitope 5. The most significant jump in possible antibody escape seen in Figure 3A corresponds to the early Omicron Nextstrain clades 21K and 21L.

**Fig 3.**
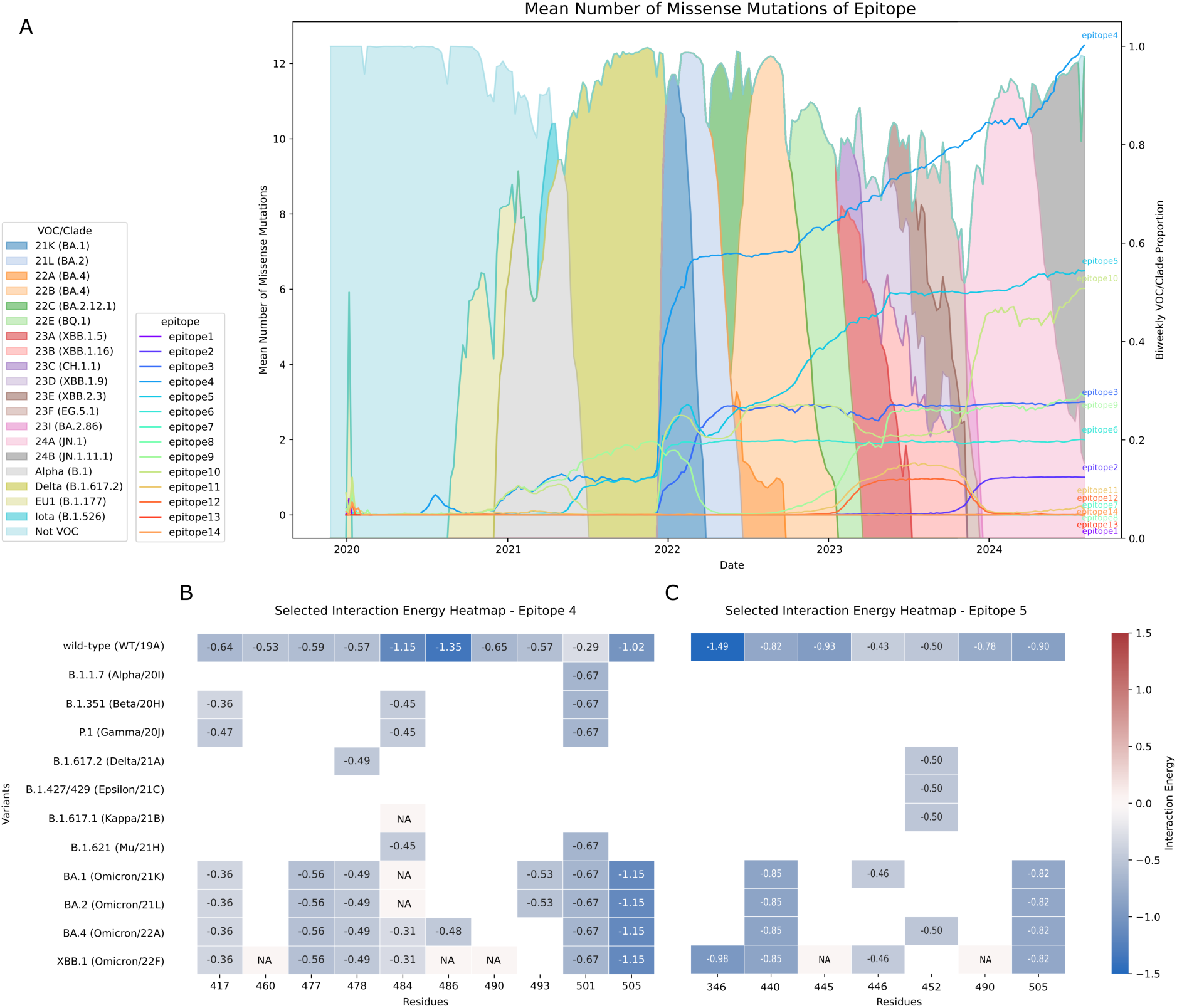
Evolution of non-synonymous mutations and interaction energies in Spike protein epitopes. (A) Timeline of mean missense mutations in 14 Spike protein epitopes across SARS-CoV-2 variants from 2020 to 2024, plotted over the bi-weekly VOC/Clade proportion. The VOCs and clades making up more than 10% of samples within each bi-weekly interval are shown. (B) Per-residue interaction energies for the mutated residues across various SARS-CoV-2 variants for epitope 4 and (C) epitope 5, with energy values (kcal/mol) calculated using Surfaces. Darker shades of blue indicate stronger favorable interactions. NA: data not available.

The Omicron VOC is the most highly mutated SARS-CoV-2 VOC to date,^53^ with 38 Spike protein mutations. The significantly altered Spike protein results in a large increase in immune escape even when compared to Delta.^54, 55^ This jump can be observed across many epitopes, but is particularly apparent in epitopes 4, 5, 10 and 6. Following this initial jump, a number of Omicron Nextstrain subclades and Pangolin lineages have emerged, with small numbers of mutations aggregating when compared to the basal 21K/21L clades.

Among the epitopes, epitope 4 has undergone the most mutations over time, followed by epitope 5. This pattern is evident as we move into the recent JN.1 and JN.1.11.1 lineages (clades 24A and 24B), which have dominated the sampled population since the start of 2024, comprising 91.28% of analysed samples from 2024 on. JN.1, with a single mutation at L455S compared to its ancestor BA.2.86, shows a notable increase in antibody evasion, particularly against Barnes’ class 1 and 2 antibodies,^56^ equivalent to epitope 4. Using the complexes with antibodies targeting epitopes 4 and 5 in variants until the recombinant XBB.1 we can pinpoint the mutations that impacted the binding affinity for those complexes. In epitope 4, substitutions in positions 417, 484 and 486 create the largest decrease in binding affinity to antibodies. For epitope 5, the R346T mutation, evaluated for its interactions in the context of the XBB.1 variant, considerably decreases antibody recognition, in accordance to experimental evaluations.^57, 58^

While mutations in epitope 5 plateaued after the emergence of XBB sublineages, epitope 10 has shown a notable rise in mutations in 2024 (Fig 3A), with substitutions such as N211I and H245N, and deletions at positions 27 and 212, over backgrounds that already carried V213G and del69. These changes underscore the adaptive pressures on regions outside the RBD, specifically in the NTD, where epitope 10 resides. However, due to the limited availability of complexes involving antibodies targeting epitope 10, it is challenging to assess the exact effects of these mutations on immune recognition based on structural evaluations.

This shift suggests that mutations in non-RBD epitopes like epitope 10 may also contribute to immune evasion, emphasizing the importance of studying and classifying epitopes beyond the RBD. Understanding mutation patterns in epitopes that experience fewer substitutions is equally valuable. Epitopes with limited mutations may serve as stable targets for antibody binding, representing potential avenues for therapeutic and vaccine design.

In our structural analysis, differentiated antibody complexes by variant to identify potential patterns of immune recognition over the course of the pandemic in the overall interactions. A key limitation of this approach is that it does not account for the origin of the antibodies or the time when they first emerged, meaning the specific variant against which these antibodies initially arose is not considered. This limitation may disproportionately affect the evaluation of the earlier variants of concern discussed here.

In the case of the Alpha variant, we observed increased antibody recognition for epitopes 4 and 5 (Fig S4B), although epitope 5 is represented by only one vector, limiting its significance (Fig S4A). The increased antibody binding affinity to epitope 4 is associated with the N501Y mutation. For the Beta variant, while the N501Y mutation enhances binding affinity at epitope 4, the overall interaction at this epitope is affected by mutations at positions 417 and 484 (Fig 8A), leading to decreased immune recognition, as previously documented.^48, 59, 60^ Additionally, we noted increased binding affinity for epitope 9, which involves interactions with mutated positions 18 and 246, and epitope 3 (Fig S4B). Regarding the Gamma variant, we observed increased binding affinity for epitopes 3, 4, and 9 (Fig S4B), although these findings are based on very few vectors and therefore not representative (Fig S4A). For epitope 9, for example, we have one vector interacting with Fab 4A8 antibody, showing lower binding affinity than *wild-type* and stronger binding affinity than BA.2 for the same antibody, while for Fab 4-8 antibody we see Gamma interacting with stronger affinity than *wild-type*. EY6A, the only epitope 3 antibody we see interacting with Gamma, and COVOX-222, the only antibody targeting epitope 4 interacting with Gamma, show similar interactions when in complex with *wild-type*, Alpha and Beta.

We also have a small subset of structures for the Mu variant, making the evaluation of the decreased binding affinity for epitope 3 less reliable (Fig S4A). Also, the Mu interaction results are based only on interactions with antibody VACW-209, and the binding affinity is similar to that of the same antibody with *wild-type* Spike and BA.1 variant. The results for the Delta variant show stronger binding affinity in epitopes 3 and 4 (Fig S4B). Considering particular antibodies enriched in the dataset, we see stronger interactions of Delta with S2L20 and VACW-209 when compared to *wild-type* and other variants.

For the Kappa variant we see, at first, a significant decrease in binding affinity for epitope 3 (Fig S4B), which is not coherent with the positions of its mutations, L452R and E484Q, located at epitope 4. Inspecting the data closely, we see that all Kappa interactions on epitope 3 are of antibody S309, the antibody with the highest representation in our dataset across different variants. Comparing all the vectors of interaction for S309, we see that one Kappa structure (PDB 7SOB), from which 3 out of the 4 vectors of interaction in epitope 3 come, is not in contact with positions 354 to 361 due to a slightly different geometry of interaction. We also see this for Omicron structures (PBD 7TM0, 7YQY), but this effect gets diluted for variants with larger subsets of structures. The effect we see, therefore, is not related to Kappa mutations. For epitope 12, all Kappa interactions are with antibody S2L20, also present in our dataset in complex with other variants. When comparing only S2L20 interactions, we see Kappa, Delta and BA.4/BA.5 — considered together due to identical mutational profiles for the Spike protein — sharing similar interactions, while Epsilon and BA.1 show decreased binding affinities.

For the BA.1 variant as a whole, there was a significant decrease in binding affinity observed for epitope 3 in general (Fig S4B), located near the mutated position 375. Interestingly, we see stronger binding affinity for BA.1 with antibody EY6A when compared to *wild-type* and other variants. The BA.4/BA.5 variant showed a significant decrease in overall strength of interactions for epitope 4, which is primarily due to the crucial mutation at position 486,^61^ along with substitutions T478K, Q498R, and S477N. Lastly, the XBB.1 variant exhibited a significant decrease in binding affinity for epitope 3 (Fig S4B), although this observation is based on only one vector of interaction of a complex with S309.

In summary, we present a comprehensive classification of 14 epitopes covering the majority of the Spike protein. This classification, based on 942 Spike-antibody complexes, provides a robust framework for the automated analysis of new complexes with a clear and replicable methodology. The evaluation of per-residue antibody recognition, performed with Surfaces, showed high correlation with escape scores calculated from DMS results. Although our per-epitope per-variant evaluation is limited by the varying compositions of the subsets, it suggests potential patterns of immune recognition and escape, particularly highlighting epitope 3 as frequently associated with significant variations in binding affinity across different variants. SPEAR was used to evaluate longitudinal trends in variants, allowing exploration of the mutations arising in variants not represented in the structural dataset, which highlighted epitopes 4 and 10 as being of interest.

### 3.2 Receptor binding affinity

The primary functional role of the Spike protein in SARS-CoV-2 cell entry involves its interaction with the receptor ACE2. Each vector of Spike-ACE2 interaction was classified into the corresponding epitope. Of the total of 209, 206 vectors are assigned to epitope 4 (Fig 1D,E).

Spike-ACE2 complexes were evaluated according to the variant they represent based on annotations from the structural dataset. Alpha,Beta, and Gamma, are consistently associated to higher ACE2 binding in experimental results,^62–64^ particularly due to the Y501-mediated increased binding.^10, 65–69^ In our results, the N501Y mutation is shown to increase binding affinity, but at the same time we see lower binding for positions 496 and 498, that compensate the 501 mutation effects (Fig 4C). The mutations on position 417, first introduced in Beta and Gamma, also appear to decrease binding according to our interaction evaluations (Fig 4C), an observation also seen through other computational^9, 10^ and experimental evaluations.^65, 67^ All variants carrying mutations on position 484 show minor effects in binding affinity in this position, also in agreement with the literature.^9, 10, 65, 67^

**Fig 4.**
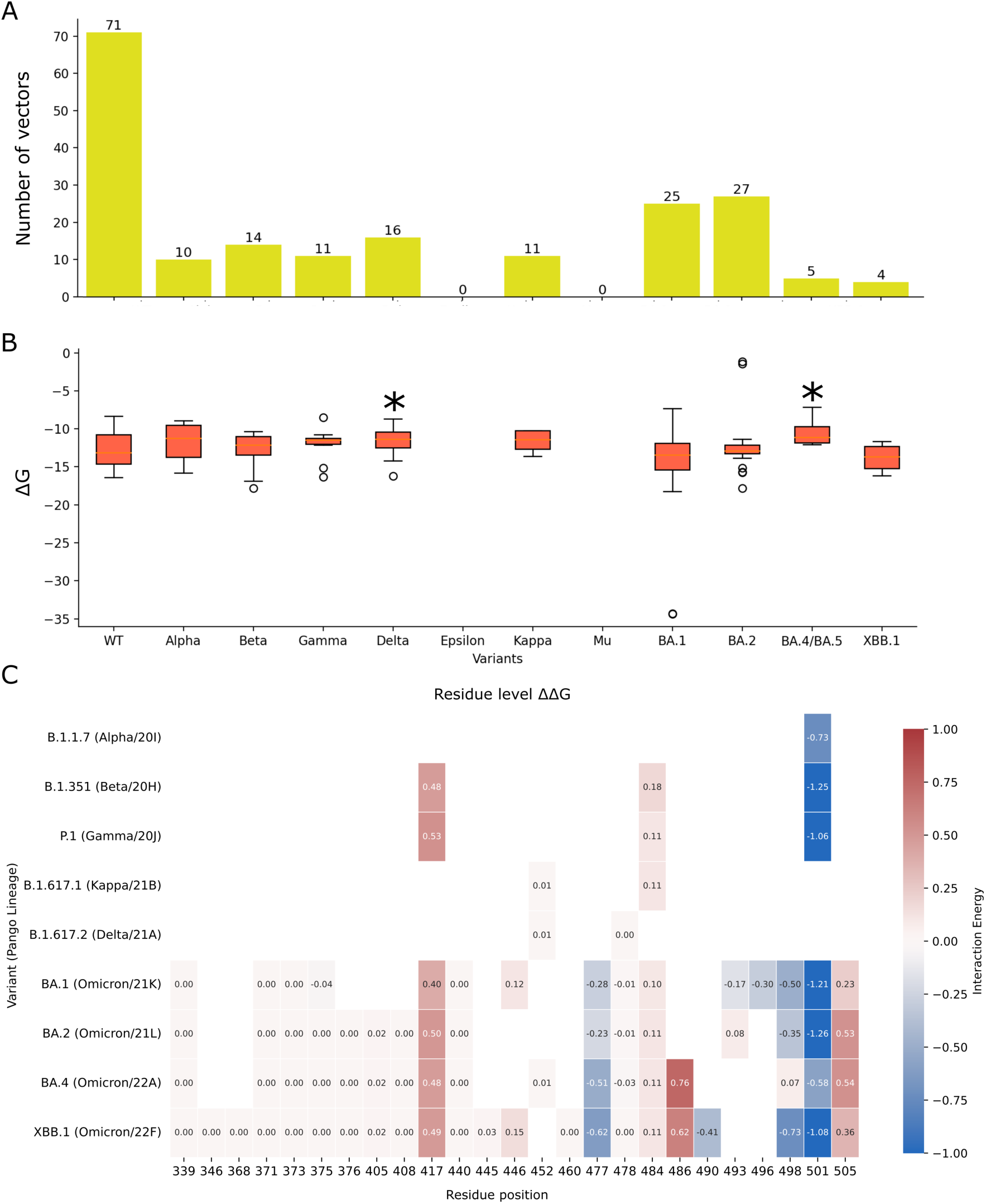
ACE2 binding calculations. (A) Composition of the dataset, showing the number of interaction vectors per variant. (B) Calculated total ACE2 interactions (kcal/mol) for structures representing each variant; ⇤*p <* 0.05. (C) Per-residue differences in average interaction between each evaluated variant and the *wild-type* ACE2 interaction for the mutated positions.

The only two variants that we were able to map as having significantly different total values of interaction compared to *wild-type* were Delta and BA.4/BA.5, both with decreased binding affinity to ACE2 (Fig 4B). Experimental data on the binding of Delta to ACE2 is conflicting, but mostly shows that it did not present much difference when compared to *wild-type*,^62, 63, 70^ which is consistent with the minor effects reported for each of its mutations individually.^10, 65^ Delta, characterized by the mutations L452R and T478K in the RBD, shows in the per-residue evaluation a decrease in binding affinity particularly in position 493, which is in close proximity to position 452 in the folded RBM. On the other hand, Kappa, that also brings the L452R substitution, as well as E484Q, shows milder effects on position 493 and slightly stronger effects in position 490, also in the neighboring region of 452. Still, it shows the smallest differences in the per-residue binding decomposition when compared to *wild-type* among all the variants analyzed. For both variants we see increased binding in position 486 and 487, that are close to the mutated K478 in Delta and the mutated Q484 in Kappa.

Mutations on position 486, present in variants BA.4/BA.5 and XBB.1, show decreased binding affinity to ACE2 (Fig 4C), in agreement with experimental evidence.^10, 65, 71, 72^ The same is true for the Y505H substitution (Fig 4C), present in Omicron subclades.^68, 73^ The S477N substitution, also part of the Omicron subclades, is associated with increased binding affinity, both in the per-residue decomposition of our analyses (Fig 4C) and in different experimental evaluations.^10, 68^ The Q498 mutation, associated to unfavorable interactions in the context of the N501Y mutation for variants Alpha, Beta and Gamma, once mutated to R498 in the Omicron subclades, shows a significant contribution to increased binding complementarity for BA.1, BA.2 and XBB.1 (Fig 4C).^69, 74^

In summary, we examined the Spike-ACE2 interaction across 209 interacting interfaces from 146 structures. From those evaluations we can characterize epitope 4 as the shared binding site between ACE2 and antibodies. Our variant-specific analysis highlighted the complex, context-specific and sometimes compensatory effects of key mutations like N501Y, K417N/T, S477N, Q498R, F486V/S, and Y505H.

### 3.3 Conformational dynamics

The conformational dynamics of the Spike protein impacts several of its functionalities. To evaluate the dynamical modulation of antibody recognition and receptor binding, we first classified the 14 epitopes based on their conformation-dependent exposure. We calculated interaction vectors between Spike chains within the trimer, in which the interacting chains could be in different conformations. Analyzing these vectors, we observe that some residues that belong to particular epitopes only become exposed when one RBD is in the open conformation and likewise, there are residues that only became exposed when two chains of Spike in the open state interact. To allow for visual comparison, we map these two cases onto the same reference structure of Spike used for all representations in Figure 1 (with one open RBD) (Fig 1G). Calculating the distances between the position of the residues that are exposed when at least one RBD is open (in green in 1G), to the position of epitope centers (Fig 1B), we can see that the exposure of epitope 4 is contingent upon at least one RBD being in the open conformation (Fig 1F), consistent with the observation that epitope 4 includes the RBM. Figure 1G also shows the residues that only became exposed when multiple RBDs are open (in purple). As can be seen in 1G, some of these in a darker shade of purple (see also the available PyMOL session, link in Data Availability) are actually occluded and would only be exposed if they were mapped onto the homotrimeric Spike structure with two RBD in the open conformation. These residues are mostly part of epitope 3 (Fig 1F). The conformation-dependent exposure of epitope 3 can be visualized in Figure S2E, in which the alignment of the interacting nanobody 17F6 (PDB 7FBJ) to the reference one-RBD-up Spike structure creates overlaps, that would prevent this binding site to exist due to steric clashes.

We selected pairs of structures representing the closed conformation and the one-RBD-open conformation, to allow us to calculate the occupancy of each of these two states for different variants (Fig 5A). This evaluation is important to estimate the exposure of these epitopes in each variant represented in our dataset of structures. We calculated the open state occupancy for each pair of structures (Fig 5B) employing the same methodology previously used to evaluate modeled single mutations,^13^ with an updated parameterization of the Markov chain (see Methods). *Wild-type* and Delta are the two groups with the most pairs of structures; 64 and 72, respectively. The high number of structures, carrying many structural variations, brings along a high dispersion of results. It is important to notice that many publications focused on experimentally investigating conformational states for different Spike variants describe sub-states and sub-populations^43, 75, 76^ that we did not consider in the evaluation; for the purpose of comparing variants, the conformational states to be compared need to be equivalent. For that reason, these sub-states are comprised on a more simplistic evaluation of closed and open, possibly contributing as well to data dispersion. From the previously modeled single mutations, we also present the Vibrational Difference Scores (VDS), a proxy for occupancy based on gain or loss of flexibility in different conformational states, for the mutations that constitute each of the variants in an attempt to interpret the reasons for occupancy shifts evaluating the effects of single substitutions (Fig 5C).

**Fig 5.**
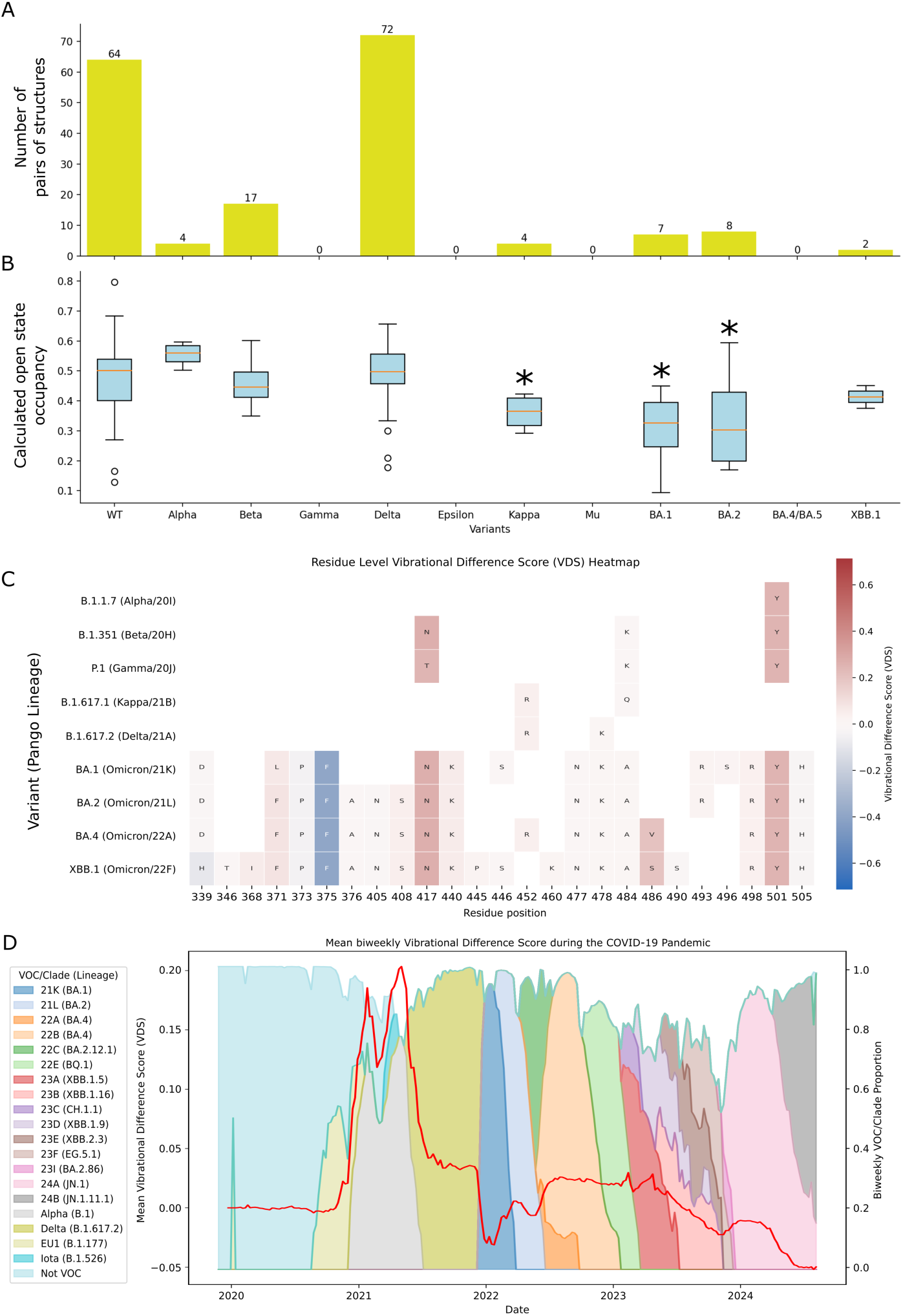
Occupancy calculations. (A) Dataset composition with the number of pairs of structures per variant. (B) Calculated open state occupancy for the pairs of structures representing each variant, ⇤*p <* 0.05. (C) Vibrational difference score (VDS) values for the mutations that constitute each variant (data from Teruel *et al.* 2021^13^), and (D) the mean bi-weekly VDS evolution during the COVID-19 pandemic, plotted over the bi-weekly VOC/Clade proportion. The VOCs and clades making up more than 10% of samples within each bi-weekly interval are shown.

Based on the distribution of points representing pairs of *wild-type* Spike structures, we do not observe a significant difference in the open state occupancy of the Alpha variant. The average calculated open state occupancy, however, is higher than *wild-type*. This result aligns with previous calculations for modeled Alpha structures,^77^ as well as with the sharp increase in sample level VDS, derived through SPEAR, corresponding to the emergence of Alpha VOC as seen in Figure 5, due to the N501Y mutation increasing the flexibility of the closed Spike and the rigidity of the open state.^13^ This observation is consistent with experimental measurements.^75^

Even with the very high dispersion of the *wild-type* data, used to determine significance, a significant reduction in the open state occupancy of Kappa, BA.1 and BA.2 variants is seen. The Omicron subclades contain the important S375F substitution,^73^ that presents a strongly negative VDS (−0.39 VDS, Fig 5C), meaning it stabilizes the closed conformation according to our previous evaluations.^13^ This observation is reflected in the longitudinal analysis of VDS (Fig 5D), in which a drop can also been seen with the onset of Clade 21K (BA.1) Omicron wave, driven by the acquisition of S375F mutation (Fig 5C). Experimental results also show a stabilized closed conformation in Omicron subclades,^40, 72, 76^ and pinpoint the role of S375F in this functional shift.^78^

We do not see in Figure 5B the same pattern of decreased open state occupancy for XBB.1, resulted from recombination between two lineages of BA.2^79^ — we see, instead, fluctuations in the calculated open state occupancies of different variants. One extra substitution that characterizes XBB.1 and was associated with increased open occupancy as a modeled single mutant is F486S (Fig 5C).^13^

Using longitudinal analysis of mutations affecting VDS, the trends observed in structures demonstrated in Figure 5B can be extended to the latest circulating variants. Here, it’s relevant to highlight substitutions L455S, L455F, and F456L, part of the JN.1-derived subvariants SLip and FLiRT, that present negative VDS,^13^ possibly contributing to the trend of stability of the closed conformation,^80^ as well as to immune escape for antibodies targeting conformation-dependent epitopes.

To summarize, we mapped epitopes 3 and 4 as conformation-specific epitopes by comparing the positions of the 14 predetermined epitopes with that of residues that exhibit conformation-dependent exposure. Our per-variant occupancy calculations suggest an initial increase in open state occupancy, primarily driven by the dynamical effects of the N501Y mutation (Fig 5C), followed by a decrease in open state occupancy in the Omicron subclades, largely due to the effects of the S375F mutation (Fig 5C), and to substitutions in positions 455 and 456 for SLip and FLiRT.

### 3.4 Glycan coating

Filtering the structures for the presence of glycans shows that the vast majority are glycosylated (Table 3). This extensive glycan coating covers various parts of the protein (Fig 1C, G). Notably, we do not observe significant glycan coverage around the RBM, likely due to natural selection favoring receptor binding — an observation in accordance to previous studies.^81^ To ensure that protein binding near the RBM does not bias this observation, we defined subsets of structures based on the presence of other macromolecules and specifically evaluated the glycan interactions of the unbound Spike (Table 3). This analysis confirms that the lack of glycosylation in the RBM region is an inherent feature of the structure and not influenced by external binding partners (Fig 1I).

**Table 3.** Composition of the dataset of structures regarding the presence and interactions of glycans.

| Subset | Number of structures | Number of structures containing glycans (percentage) | Number of structures with non-null interactions between glycans and Spike (percentage) | Number of structures with glycans co-interacting with more than one chain of Spike (percentage) | Number of structures with non-null interactions between glycans and the interacting protein (percentage) | Number of structures with glycans co-interacting with Spike and its interacting protein (percentage) |
| --- | --- | --- | --- | --- | --- | --- |
| Spike + antibody | 942 | 723 (76.8%) | 721 (76.5%) | 101 (10.7%) | 201 (21.3%) | 184 (19.5%) |
| Spike + ACE2 | 146 | 118 (78.6%) | 111 (76.0%) | 39 (26.7%) | 107 (73.3%) | 16 (11.0%) |
| Unbound Spike | 338 | 269 (80.8%) | 269 (80.8%) | 261 (77.2%) | - | - |

Similar glycan coating patterns appear in the other two subsets — structures of Spike in complex with antibodies and with the receptor ACE2 (Fig S5A, B). In the antibody complexes, glycans coat key immune recognition epitopes (Fig S5C, D), raising the possibility that glycans contribute to immune escape. In glycosylated structures bound to ACE2, glycans interact with residues near the peripheral RBM region (Fig S5E, F), though the glycans attach to residues in neighboring Spike chains rather than within the RBM itself, co-interacting with the closest RBD (Fig S5G). We also observe glycans co-interacting with Spike and antibody chains (Fig S5G). Their presence may not be obligatory, as some antibody complexes targeting the same epitope exhibit high surface complementarity without glycans (Fig S5I). Examples like these prompted us to search the entire dataset for co-interacting glycans to further investigate their role in immune recognition (Table 3).

Investigating the effects of the most common co-interacting glycan, we identified 203 interaction vectors influenced by the glycan linked to asparagine 343 (ASN343) (Fig 6A). In the main interacting Spike chain, this glycan increases the binding surface area by introducing new interacting residues, resulting in an average 12.65% increase in binding affinity, with the largest gain reaching 4.56 kcal/mol (Fig 6D, E, F).

**Fig 6.**
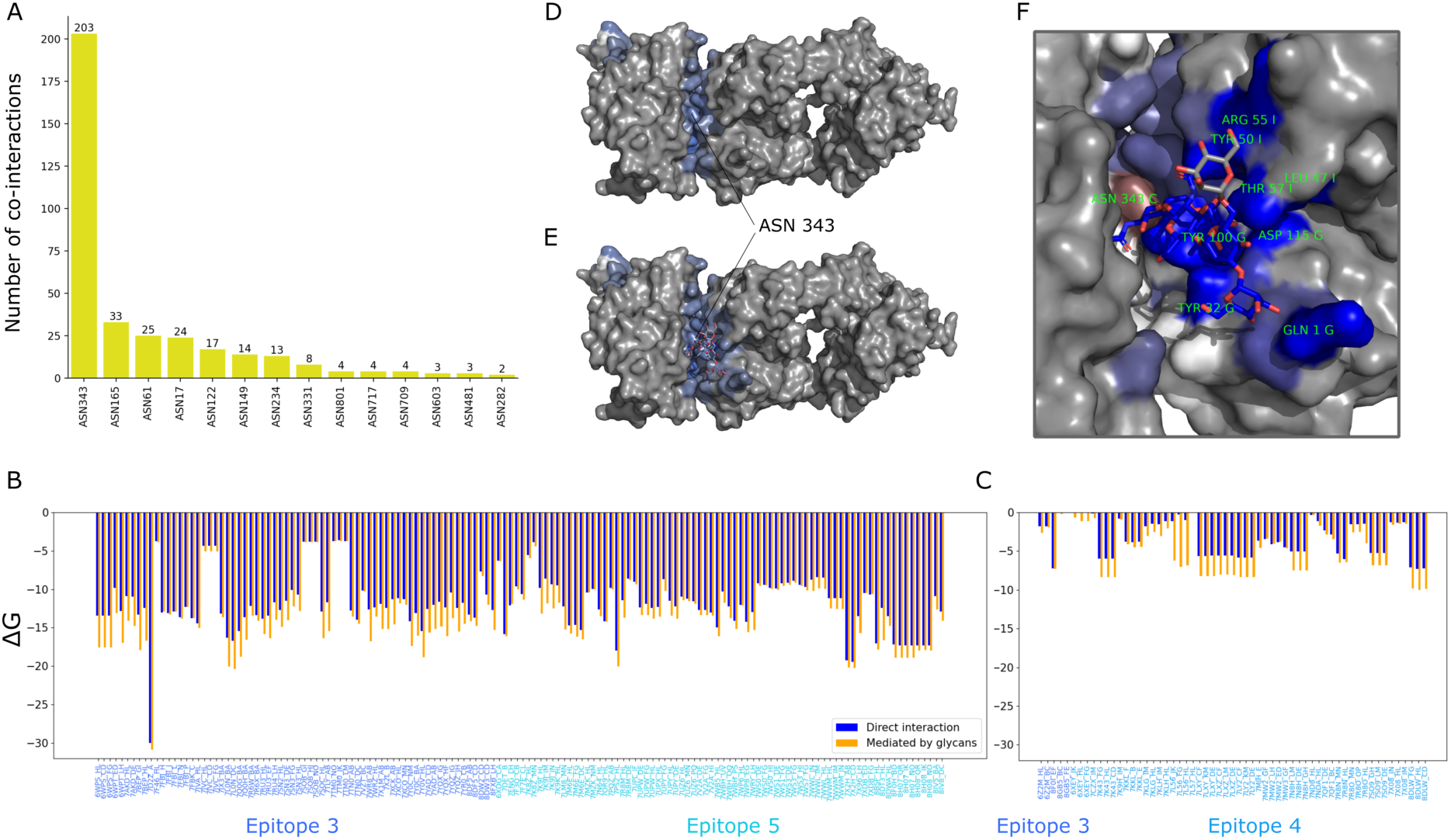
Characterization of glycan-mediated antibody binding. (A) Number of co-interactions per glycan-linked residue. (B) Values of binding affinity (Δ*G*, kcal/mol) comparing direct interaction (blue) to interaction mediated by ASN343-linked glycans (orange) for co-interactions in which the glycan mediated binding of antibody chains to its main interacting Spike chain, or (C) a neighboring Spike chain. (D) Direct interaction between Spike RBD and the S309 antibody (PDB 7BEP), (E) and the same interaction mediated by ASN343-linked glycan, showing an increased binding site. (F) Close-up of the delta between results, showing gain of interactions for most antibody residues, namely GLN1, TYR32, TYR100, and ASP115 from S309 heavy chain and LEU471, TYR501, and ARG551 from S309 light chain, and a loss of direct interaction of Spike’s ASN343.

Considering the specific epitopes for each interaction, ASN343 glycosylation mediates interactions in epitopes 3 and 5. The evaluation of glycan mediation per epitope reveals an average interaction gain of 2.00 kcal/mol and 1.07 kcal/mol, as predicted by Surfaces, representing a 16.50% and 9.21% increase in affinity, respectively (Fig 6B). Beyond enhancing interactions with the main interacting Spike chain, the ASN343-linked glycan significantly increases interactions with neighboring Spike chains, especially for antibodies targeting epitope 4. This effect is pronounced in structures like PDB 7L56, where glycan mediation greatly multiplies interactions (Fig 6C). Including these interactions, the largest binding affinity gain reaches 6.77 kcal/mol.

We also observed interactions being mediated by glycans linked to other asparagine residues (Fig S6A). Noteworthy examples include strengthened interactions between neighboring chains, mediated by glycans linked to ASN165 and ASN122 (Fig S6B), glycan-dependent interactions of antibody 2G12 targeting epitope 8 via glycans linked to ASN801, ASN717, and ANS709 (Fig S6C),^82, 83^ and enhanced interactions in antibodies targeting epitope 13 due to glycans linked to ASN61, ASN282, and ANS603 (Fig S6D).

All co-interacting glycans linked to Spike residues mediate antibody interactions. Some structures reveal glycans co-interacting with both Spike and ACE2, all linked to ACE2 residues (Fig S7A, C) and playing important roles in ACE2 recognition.^84–86^ We also observed a few antibody complexes where glycans link to residues within the antibody chains, though these cases are rare (Fig S7B, D).

Beyond mediating antibody interactions, we investigated whether glycans can mask specific epitopes. Some residues, such as ASN1074, ASN1098, ASN1134, and ANS1158 in epitope 1, are never glycosylated in the context of antibody binding to the same epitope — when evaluating each structure, the presence of these glycans and of antibodies bound to epitope 1 appears to be mutually exclusive, suggesting that the recognition of epitope 1 is masked by the glycan shield. Glycans only link to these residues in structures without antibody in complex or where the antibodies target different epitopes.

While selecting structures in our dataset in which there are glycans co-interacting with more than one protein unit, we identified many structures where glycans co-interact with more than one Spike protein chain. To assess their role in mediating conformational dynamics, we examined how frequently these interactions occur, along with the interaction values between the linked chain and the co-interacting chain, across different Spike conformational states (Fig 7A, B, C, D). Glycans linked to ANS709 and ASN1074 often co-interact with neighboring Spike chains, but the distribution of binding affinities remains unchanged across conformational states, as both positions are distant from the RBD. However, for ASN234-linked glycans, we observe conformation-specific results, with significantly stronger affinity for the co-interacting chain when both chains are in the down conformation (Fig 7B, F, G). Glycans linked to ASN343 and ASN370 also show conformation-specific interactions, occurring only when both chains are in the down conformation (Fig 7B, H, I). ASN234, ASN343 and ASN370-linked glycans had previously been described for their role modulating the opening of the RBD;^87–90^ the same is true fot ASN165-linked glycans, but our interactions results did not show conformation-specific changes in binding affinity. Based on the same evaluation of enriched results, the ASN1158 position can also be highlighted (Fig 7E), but the absence of co-interactions between two Spike chains in the up conformation likely stems from the low resolution of the C-terminal in many homotrimeric structures.

**Fig 7.**
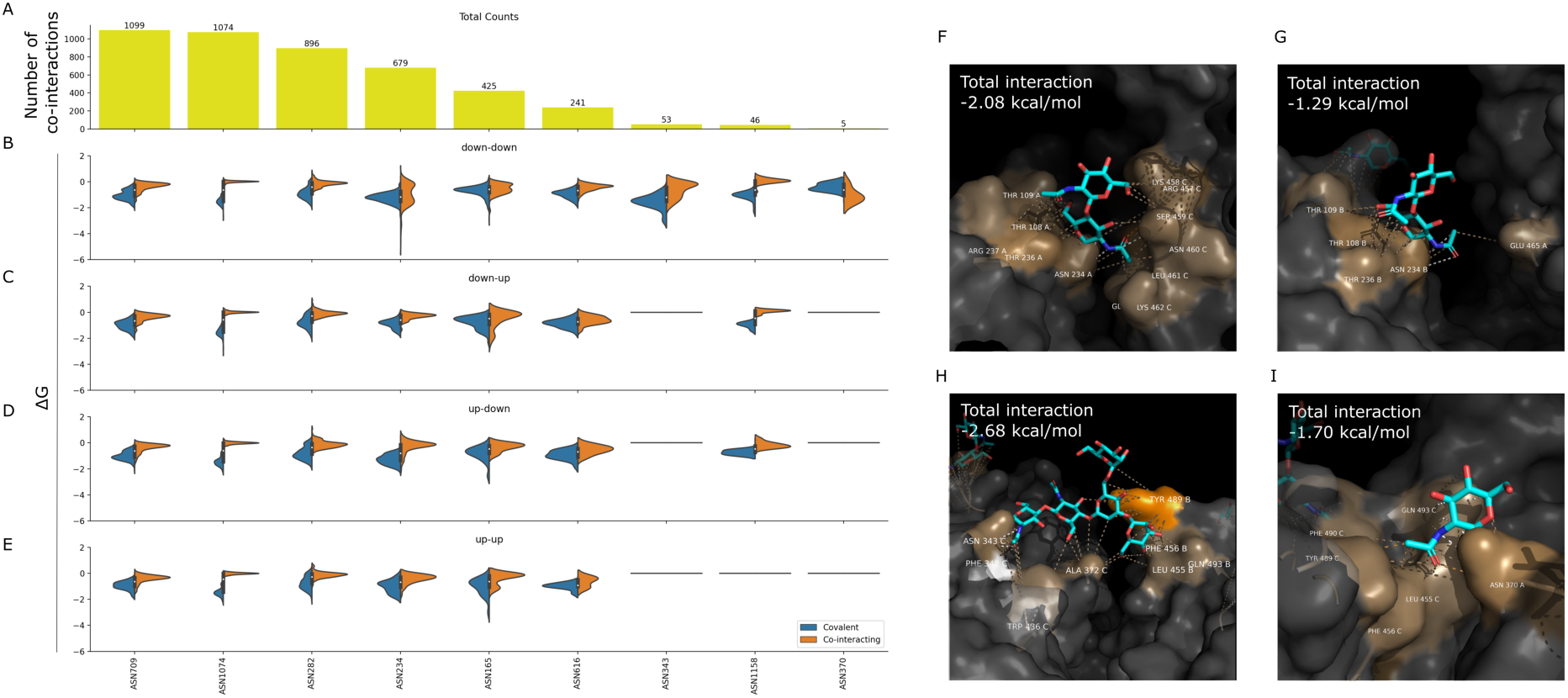
Characterization of glycan-mediated conformational stability. (A) Number of co-interacting glycans based on the Spike residue they are covalently linked to. (B) Distributions of binding affinity (Δ*G*, kcal/mol) of glycans bound to a Spike chain in the down conformation and co-interacting with a Spike chain in the down conformation, (C) bound to a Spike chain in the down conformation and co-interacting with a Spike chain in the up conformation, (D) bound to a Spike chain in the up conformation and co-interacting with a Spike chain in the down conformation, and (E) bound to a Spike chain in the up conformation and co-interacting with a Spike chain in the up conformation, with the distribution of total affinity for the chain they are covalently bound to (blue) and the co-interacting chain (orange). For glycans linked to residue ASN234, the affinity to the co-interacting chain is much higher for chains in the down conformation (F) then in the up conformation (G), as exemplified by two different pairs of Spike chains from the same structure (PDB 6XM3). We can also see examples of glycans (H) linked to ASN343 (PDB 8EPN), as well as (I) to ASN370 (PDB 7FCE), stabilizing the interaction between a pair of closed Spike chains.

Due to the inherent flexibility of glycans, their resolution is often limited in structural data. Since we did not manipulate the structures or reconstruct the entire glycan chains, this may have influenced our evaluations. By focusing only on the portions of glycans explicitly resolved in the structures, our analysis is inherently biased by the resolution of the data. However, the glycosylated positions we highlight with conformation-specific glycan interactions are typically evaluated within the same structure, comparing chains in different conformational states (Fig 7F, G), and are therefore subject to consistent resolution constraints. Furthermore, the most flexible regions of the glycans are less constrained due to the absence of strong interactions, meaning these regions would not significantly contribute to our interaction evaluations.

Given the functional relevance of glycans linked to specific residues, we assessed their conservation to identify potential mutations that could disrupt glycosylation and affect function indirectly. All positions selected for their potential impact on protein dynamics and antibody recognition are highly conserved according to CoV-Spectrum data^91^(Table S1), aligning with other computational studies that suggest Spike protein mutations typically occur in regions where antibody access is not obstructed by glycans.^92^ Although our occupancy calculations did not account for glycan-mediated interactions between Spike chains, this exclusion does not affect our conclusions, as our analysis focused on vibrational entropy differences relative to *wild-type* structures, which similarly excluded glycan influence. However, experimental evidence points to varying glycosylation patterns across Spike variants, which may influence the glycan-mediated functionalities described here.^93–95^ These effects are primarily attributed to glycan composition, which we did not evaluate in this study. The gain of N-glycosylated sites in particular lineages,^94, 96^ such as at ASN354,^97^ has also been reported. These additions, while having the potential to affect our conclusions, cannot be assessed within our methodology due to the absence of structures with glycans bound to these sites of interest in our dataset. Additionally, the glycosylation patterns of Spike can be affected by the specific cell types in which it is expressed.^98^ As this data is not readily available programmatically, it is possible that the patterns of glycan interactions observed could not represent those in vivo. However, to the extent that such an experimental artifact may be averaged out, the fact that we utilized approximately 1,000 Spike proteins for the analysis of glycosylation, may restrict the extent of this problem.

### 3.5 Enthalpic trade-off hypothesis: Receptor binding affinity and antibody recognition

The enthalpic trade-off hypothesis is based on the underlying assumption, as previously proposed,^32, 77, 99^ that mutations in the same surface of the Spike protein will affect the binding of all proteins that interact with this epitope. Specifically, we analyzed antibody interactions at epitope 4, which shares its binding interface with the ACE2 receptor (Fig 1D). Given the dataset composition, we evaluated the interplay between antibody and ACE2 binding for the Alpha, Beta, Gamma, Delta, BA.1, BA.2, and BA.4/BA.5 variants. We compared ACE2 and antibody binding using two approaches: A per-residue decomposition of binding affinity (Fig 8A), and an analysis of total per-variant interactions (Fig 8B).

**Fig 8.**
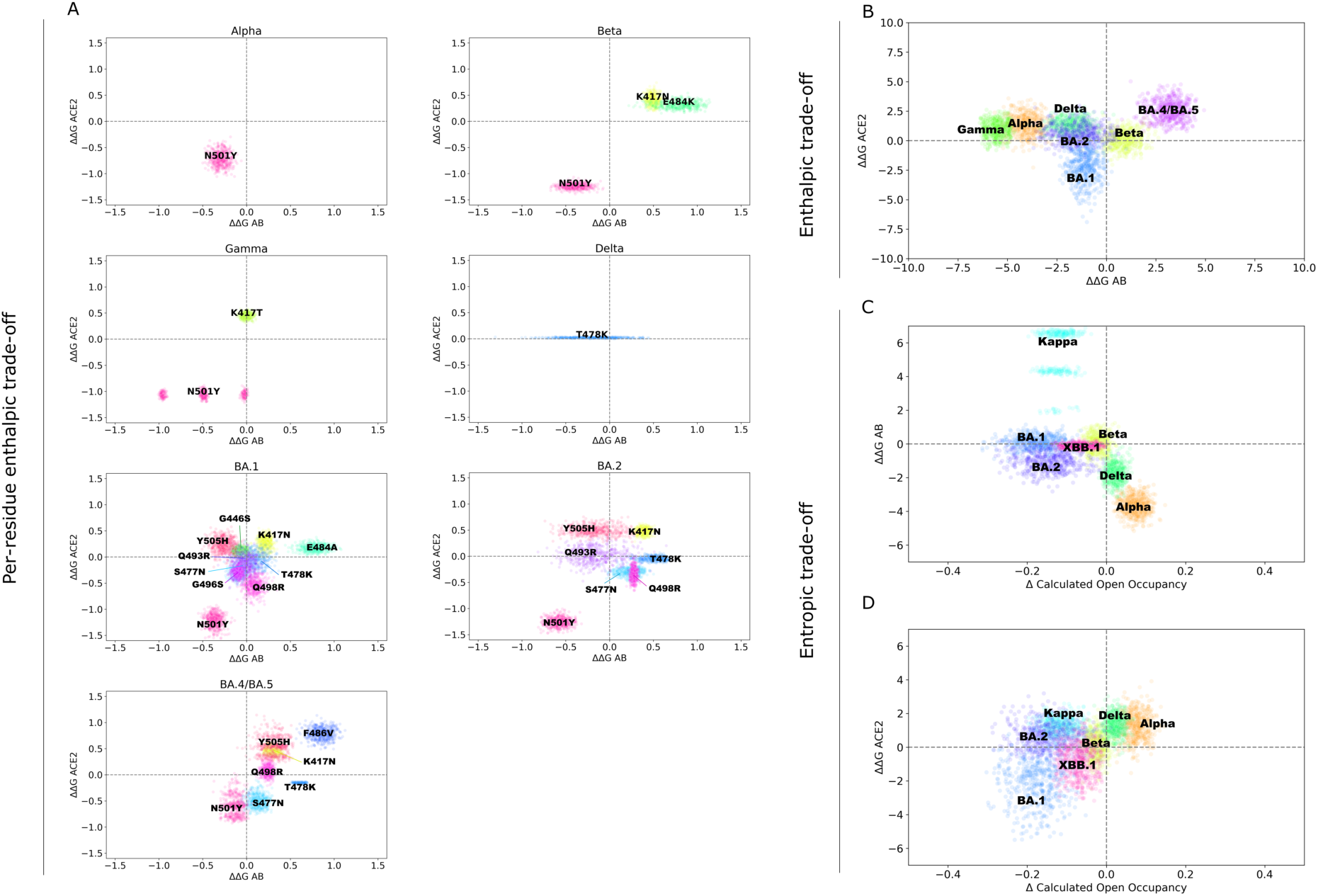
Comparative metrics of ACE2 binding affinity (ΔΔ*G* ACE2, kcal/mol), antibody recognition (ΔΔ*G* AB, kcal/mol) and conformational dynamics for Spike variants (Δ Calculated Open Occupancy, %). (A) Results of binding affinity with ACE2 and with antibodies for each mutated residue in the RBM of structures representing variants Alpha, Beta, Gamma, Delta, BA.1, BA.2 and BA.4/BA.5, and (B) full interaction results for the same structures. (C) Results of open occupancy calculations based on structures of variants Alpha, Beta, Delta, Kappa, BA.1, BA.2 and XBB.1 against the full interactions of the same variants with the receptor ACE2, or (D) with antibodies targeting conformation-specific epitopes.

For variants like Alpha and Beta, per-residue evaluations show that mutations in epitope 4 affect both ACE2 and antibodies targeting the RBM in similar ways, either increasing both interactions (N501Y) or decreasing both (K417N, E484K), aligning with the enthalpic trade-off hypothesis. For Gamma, the K417T mutation shows a more subtle immune escape effect, slightly increasing or decreasing binding affinity to antibodies, which may reflect the dataset limitations for Gamma (see Antibody recognition). The only Delta mutation detected in interactions with both ACE2 and antibodies targeting epitope 4 occurs at position 478, showing no effects on ACE2 binding but variable effects on antibody recognition.

Omicron subclades carry many mutations in the RBM. In BA.1, the effects of mutations at positions 501, 484, and 417 resemble those in Beta, while new substitutions show milder effects. T478K causes a small decrease in antibody recognition and a slight increase in ACE2 binding, similar to Q493R (Fig 8A). In the context of BA.1, Q498R also presents this effect, with a more prominent increase in ACE2 binding. S477N and G496S increase both ACE2 and antibody binding slightly. Y505H and G446S cause a minor increase in antibody binding and a decrease in ACE2 binding (Fig 8A). In BA.2, T478K, Q498R, and S477N shift toward less favorable antibody binding (Fig 8A). For BA.4/BA.5, Y505H and N501Y mutations show reduced antibody recognition, marking the first time N501Y does not strongly favor antibody binding (Fig 8A). This suggests a possible epistatic effect for Omicron subclades, shifting the per-residue binding contribution of residues 501, 505, 477, and 478 toward decreased immune recognition. BA.4/BA.5 also introduces the F486V mutation, significantly reducing binding to both RBM-targeting antibodies and ACE2. In the full interaction analysis (Fig 8B), BA.1 shows the most significant dual effect, increasing binding for both antibodies and ACE2, while BA.4/BA.5 decreases both interactions.

Considering all vectors of ACE2 interaction, we can search for the vectors of antibody interactions that most closely resemble interactions with the receptor. These are of antibodies P2B-1A1 (PDB 7CZP), Beta-29 (PDB 7PS2), and CTC-445.2 inhibitor (PDB 7KL9), that may increase the enthalpic trade-off observation and force a diminished ACE2 binding capacity for mutations selected for their immune escape effects. The binding similarity to ACE2 possibly makes these antibodies valuable in guiding the design of new therapeutic strategies.

In summary, our analysis shows a number of mutations supporting the enthalpic trade-off hypothesis, notably mutations N501Y, K417N, E484K/A, and F486V.

### 3.6 Entropic trade-off hypothesis: Conformational dynamics and RBM interactions

Whereas the enthalpic trade-off hypothesis has been previously suggested, the combined analysis of interaction enthalpies and their effect on dynamics allow us to introduce the concept of entropic trade-off in the evolution of SARS-CoV-2, that may contribute to the evolution of any virus in which different conformational states of viral proteins are involved in the cell entry mechanism. The entropic trade-off hypothesis stems from the relationship between conformational dynamics and receptor/antibody binding (Fig 8C, D). Studies have linked the closed conformation of the SARS-CoV-2 Spike protein to immune escape,^100^ a phenomenon particularly relevant for variants like BA.1 and BA.2.^14^ These variants prioritize a closed conformation as a strategy to evade immune recognition, even at the cost of reduced ACE2 binding affinity due to decreased RBM exposure.

Conformation-specific antibody binding, especially for epitopes 3 and 4, warrants particular attention, as discussed earlier (Fig 1F). Given the dataset composition, we evaluated the interplay between conformational dynamics and conformation-specific binding for Alpha, Beta, Delta, Kappa, BA.1, BA.2, and XBB.1 variants. The Omicron subclades (BA.1, BA.2, and the recombinant XBB.1) demonstrate a trend toward favoring the closed conformation, while the effects on antibody binding remain mild or even increase in the case of BA.1 (Fig 8C). This observation suggests that immune escape in these variants stems from their conformational preference for the closed state, rather than a reduction in binding affinity at the interface. When analyzing ACE2 binding and conformational dynamics, we observe increased ACE2 binding affinity in BA.1 and XBB.1, though effective binding may be compensated due to the lower occupancy of the open state.

## 4 Conclusion

The vast structural data on the SARS-CoV-2 Spike protein offers unprecedented opportunities for in-depth analysis using high-throughput computational methods. Our study shows how such an analysis can provide new insights into functional aspects of the Spike protein, particularly through a data-driven epitope classification system. By applying this approach, we categorized epitopes and developed a straightforward method for sorting interacting complexes within epitope clusters. We identified 14 distinct epitopes based on their conformational specificity, ACE2 binding, and glycan coverage, providing a detailed understanding of binding across different Spike protein domains. Our per-residue evaluations of antibody binding show high accuracy compared with experimentally determined immune escape scores for antibodies targeting the RBD, reinforcing confidence in our conclusions regarding less-studied domains also represented in our epitopes and providing a method for the evaluation of future strains or for different viruses. A longitudinal analysis of mutations within each epitope reveals that mutations in the NTD merit consideration for their potential effects on immune recognition in the latest variants. This analysis also highlights opportunities for antibody development and therapeutic alternatives targeting well-conserved epitopes. This new epitope classification and mapping facilitates a focused analysis of immune escape by narrowing it to the relevant subset of antibody structures associated with specific epitopes. For example, when studying ACE2 binding, by restricting the analysis to those antibody structures that have interactions with the RBM.

Our findings emphasize the importance of understanding how various functional aspects of the Spike protein interrelate, including its conformational dynamics, ACE2 receptor binding, antibody recognition, and glycan coating. While previous studies often evaluated these features separately, our comprehensive analysis reveals their interdependence and how they collectively influence the function and evolution of the Spkike protein and consequently of the SARS-CoV-2 virus. By evaluating these aspects per variant, we connected them within our trade-off hypothesis and conducted a per-residue analysis to explain the success of specific substitutions as they first emerged. We also explored the epistatic effects of mutations in different variant contexts, such as the varying impacts on antibody recognition from mutations at positions 477, 478, 501, and 505 in various Omicron subclades. Regarding dynamical effects, we identified N501Y as a key mutation increasing open state occupancy in early variants of concern, such as Alpha, Beta, and Gamma, while S375F in Omicron subclades indirectly facilitates immune escape by reducing open state occupancy. While the conformational shift toward the closed state of Spike has been suggested as a mechanism for immune escape, we introduce a methodology based on biophysical concepts to quantify these distinct functional effects across mutations and variants. We hope that the now clearly defined concepts of enthalpic and entropic trade-off will help characterize new mutations and understand the evolution of future variants of SARS-CoV-2 as well other viruses. Additionally, structural evaluations highlight the role of glycosylation, particularly ASN343-linked glycans in mediating antibody recognition and ASN234-linked glycans in modulating dynamics.

Understanding these connections is essential for gaining deeper insights into the evolutionary pressures acting on the Spike protein. The methods and findings presented in this study lay a solid foundation for future research into the structural biology of the Spike protein, offering new avenues for exploring its role in the adaptability and pathogenicity of SARS-CoV-2. Additionally, the methodologies we introduce can be used for the analysis of other large structural datasets, a need that is becoming increasingly relevant with the growing number of available structures, both by the popularization of structural biology techniques for structure determination and the ongoing revolution in protein modeling.

## 5 Data availability

The data supporting the findings of this study are publicly available at https://github.com/ `nataliateruel/Comprehensive_review_Spike`. This repository includes all PyMOL sessions used to generate the images, the raw data for interaction evaluations and dynamical assessments, the mutations per epitope per variant from the longitudinal studies, as well as Python scripts for classifying complexes among the 14 epitopes described.

## Supporting information

Supplementary Tables and Images

## Acknowledgments

RJN is a member of the Quebec network for the study of protein structure, function and engineering (PROTEO). MC and MB are supported by Research England’s Expanding Excellence in England (E3) Fund. The authors would like to thank Dr. Lok Yin Roy Wong at Rutgers University, Dr. Julie Hussin and Dr. Rikard Blunck at Université de Montréal and Dr. Sophie Gobeil at Université Laval for critical reading of the manuscript.

## References

1 Fan Wu, Su Zhao, Bin Yu, Yan-Mei Chen, Wen Wang, Zhi-Gang Song, Yi Hu, Zhao-Wu Tao, Jun-Hua Tian, Yuan-Yuan Pei, et al. A new coronavirus associated with human respiratory disease in china. Nature, 579(7798):265–269, 2020.

2 Roujian Lu, Xiang Zhao, Juan Li, Peihua Niu, Bo Yang, Honglong Wu, Wenling Wang, Hao Song, Baoying Huang, Na Zhu, et al. Genomic characterisation and epidemiology of 2019 novel coronavirus: implications for virus origins and receptor binding. The lancet, 395(10224):565–574, 2020.

3 Peng Zhou, Xing-Lou Yang, Xian-Guang Wang, Ben Hu, Lei Zhang, Wei Zhang, Hao-Rui Si, Yan Zhu, Bei Li, Chao-Lin Huang, et al. A pneumonia outbreak associated with a new coronavirus of probable bat origin. nature, 579(7798):270–273, 2020.

4 Wenhui Li, Michael J Moore, Natalya Vasilieva, Jianhua Sui, Swee Kee Wong, Michael A Berne, Mohan Somasundaran, John L Sullivan, Katherine Luzuriaga, Thomas C Greenough, et al. Angiotensin-converting enzyme 2 is a functional receptor for the sars coronavirus. Nature, 426(6965):450–454, 2003.

5 Xiuyuan Ou, Yan Liu, Xiaobo Lei, Pei Li, Dan Mi, Lili Ren, Li Guo, Ruixuan Guo, Ting Chen, Jiaxin Hu, et al. Characterization of spike glycoprotein of sars-cov-2 on virus entry and its immune cross-reactivity with sars-cov. Nature communications, 11(1):1620, 2020.

6 Alexandra C Walls, Young-Jun Park, M Alejandra Tortorici, Abigail Wall, Andrew T McGuire, and David Veesler. Structure, function, and antigenicity of the sars-cov-2 spike glycoprotein. Cell, 181(2):281–292, 2020.

7 Yasunori Watanabe, Joel D Allen, Daniel Wrapp, Jason S McLellan, and Max Crispin. Sitespecific glycan analysis of the sars-cov-2 spike. Science, 369(6501):330–333, 2020.

8 Cong Xu, Wenyu Han, and Yao Cong. Cryo-em and cryo-et of the spike, virion, and antibody neutralization of sars-cov-2 and vocs. Current Opinion in Structural Biology, 82:102664, 2023.

9 Bahaa Jawad, Puja Adhikari, Rudolf Podgornik, and Wai-Yim Ching. Key interacting residues between rbd of sars-cov-2 and ace2 receptor: combination of molecular dynamics simulation and density functional calculation. Journal of chemical information and modeling, 61(9):4425–4441, 2021.

10 Alina P Sergeeva, Phinikoula S Katsamba, Junzhuo Liao, Jared M Sampson, Fabiana Bahna, Seetha Mannepalli, Nicholas C Morano, Lawrence Shapiro, Richard A Friesner, and Barry Honig. Free energy perturbation calculations of mutation effects on sars-cov-2 rbd:: Ace2 binding affinity. Journal of Molecular Biology, 435(15):168187, 2023.

11 Rachael A Mansbach, Srirupa Chakraborty, Kien Nguyen, David C Montefiori, Bette Korber, and Gnana Gnanakaran. The sars-cov-2 spike variant d614g favors an open conformational state. Biophysical Journal, 120(3):298a, 2021.

12 Gennady M Verkhivker. Molecular simulations and network modeling reveal an allosteric signaling in the sars-cov-2 spike proteins. Journal of proteome research, 19(11):4587–4608, 2020.

13 Natália Teruel, Olivier Mailhot, and Rafael J Najmanovich. Modelling conformational state dynamics and its role on infection for sars-cov-2 spike protein variants. PLoS computational biology, 17(8):e1009286, 2021.

14 Valeria Calvaresi, Antoni G Wrobel, Joanna Toporowska, Dietmar Hammerschmid, Katie J Doores, Richard T Bradshaw, Ricardo B Parsons, Donald J Benton, Chloë Roustan, Eamonn Reading, et al. Structural dynamics in the evolution of sars-cov-2 spike glycoprotein. Nature Communications, 14(1):1421, 2023.

15 William T Harvey, Alessandro M Carabelli, Ben Jackson, Ravindra K Gupta, Emma C Thomson, Ewan M Harrison, Catherine Ludden, Richard Reeve, Andrew Rambaut, Sharon J Peacock, et al. SARS-CoV-2 variants, spike mutations and immune escape. Nature Reviews Microbiology, 19(7):409–424, 2021.

16 Luca Piccoli, Young-Jun Park, M Alejandra Tortorici, Nadine Czudnochowski, Alexandra C Walls, Martina Beltramello, Chiara Silacci-Fregni, Dora Pinto, Laura E Rosen, John E Bowen, et al. Mapping neutralizing and immunodominant sites on the SARS-CoV-2 spike receptor-binding domain by structure-guided high-resolution serology. Cell, 183(4):1024–1042, 2020.

17 Ellen Shrock, Eric Fujimura, Tomasz Kula, Richard T Timms, I-Hsiu Lee, Yumei Leng, Matthew L Robinson, Brandon M Sie, Mamie Z Li, Yuezhou Chen, et al. Viral epitope profiling of COVID-19 patients reveals cross-reactivity and correlates of severity. Science, 370(6520):eabd4250, 2020.

18 Christopher O Barnes, Claudia A Jette, Morgan E Abernathy, Kim-Marie A Dam, Shannon R Esswein, Harry B Gristick, Andrey G Malyutin, Naima G Sharaf, Kathryn E Huey-Tubman, Yu E Lee, et al. Sars-cov-2 neutralizing antibody structures inform therapeutic strategies. Nature, 588(7839):682–687, 2020.

19 Christopher O Barnes, Anthony P West, Kathryn E Huey-Tubman, Magnus AG Hoffmann, Naima G Sharaf, Pauline R Hoffman, Nicholas Koranda, Harry B Gristick, Christian Gaebler, Frauke Muecksch, et al. Structures of human antibodies bound to sars-cov-2 spike reveal common epitopes and recurrent features of antibodies. Cell, 182(4):828–842, 2020.

20 Ashlesha Deshpande, Bethany D Harris, Luis Martinez-Sobrido, James J Kobie, and Mark R Walter. Epitope classification and rbd binding properties of neutralizing antibodies against sars-cov-2 variants of concern. Frontiers in immunology, 12:691715, 2021.

21 Kathryn M Hastie, Haoyang Li, Daniel Bedinger, Sharon L Schendel, S Moses Dennison, Kan Li, Vamseedhar Rayaprolu, Xiaoying Yu, Colin Mann, Michelle Zandonatti, et al. Defining variant-resistant epitopes targeted by SARS-CoV-2 antibodies: A global consortium study. Science, 374(6566):472–478, 2021.

22 Meng Yuan, Deli Huang, Chang-Chun D Lee, Nicholas C Wu, Abigail M Jackson, Xueyong Zhu, Hejun Liu, Linghang Peng, Marit J Van Gils, Rogier W Sanders, et al. Structural and functional ramifications of antigenic drift in recent SARS-CoV-2 variants. Science, 373(6556):818–823, 2021.

23 Meghan E Garrett, Jared G Galloway, Caitlin Wolf, Jennifer K Logue, Nicholas Franko, Helen Y Chu, Frederick A Matsen IV, and Julie M Overbaugh. Comprehensive characterization of the antibody responses to sars-cov-2 spike protein finds additional vaccine-induced epitopes beyond those for mild infection. Elife, 11:e73490, 2022.

24 Yanjia Chen, Xiaoyu Zhao, Hao Zhou, Huanzhang Zhu, Shibo Jiang, and Pengfei Wang. Broadly neutralizing antibodies to SARS-CoV-2 and other human coronaviruses. Nature Reviews Immunology, 23(3):189–199, 2023.

25 Arnaud N’Guessan, Senthilkumar Kailasam, Fatima Mostefai, Raphaël Poujol, Jean-Christophe Grenier, Nailya Ismailova, Paola Contini, Raffaele De Palma, Carsten Haber, Volker Stadler, et al. Selection for immune evasion in sars-cov-2 revealed by high-resolution epitope mapping and sequence analysis. Iscience, 26(8), 2023.

26 Ayijiang Yisimayi, Weiliang Song, Jing Wang, Fanchong Jian, Yuanling Yu, Xiaosu Chen, Yanli Xu, Sijie Yang, Xiao Niu, Tianhe Xiao, et al. Repeated omicron exposures override ancestral sars-cov-2 immune imprinting. Nature, 625(7993):148–156, 2024.

27 Rajeshwer S Sankhala, Vincent Dussupt, Wei-Hung Chen, Hongjun Bai, Elizabeth J Martinez, Jaime L Jensen, Phyllis A Rees, Agnes Hajduczki, William C Chang, Misook Choe, et al. Antibody targeting of conserved sites of vulnerability on the sars-cov-2 spike receptor-binding domain. Structure, 32(2):131–147, 2024.

28 Yuelong Shu and John McCauley. Gisaid: Global initiative on sharing all influenza data– from vision to reality. Eurosurveillance, 22(13):30494, 2017.

29 Stefan Elbe and Gemma Buckland-Merrett. Data, disease and diplomacy: Gisaid’s innovative contribution to global health. Global challenges, 1(1):33–46, 2017.

30 James Hadfield, Colin Megill, Sidney M Bell, John Huddleston, Barney Potter, Charlton Callender, Pavel Sagulenko, Trevor Bedford, and Richard A Neher. Nextstrain: real-time tracking of pathogen evolution. Bioinformatics, 34(23):4121–4123, 2018.

31 Sonja Marjanovic, Robert J Romanelli, Gemma-Claire Ali, Brandi Leach, Margaretha Bonsu, Daniela Rodriguez-Rincon, and Tom Ling. Covid-19 genomics uk (cog-uk) consortium. Rand Health Quarterly, 9(4), 2022.

32 Marine E Bozdaganyan, Konstantin V Shaitan, Mikhail P Kirpichnikov, Olga S Sokolova, and Philipp S Orekhov. Computational analysis of mutations in the receptor-binding domain of sars-cov-2 spike and their effects on antibody binding. Viruses, 14(2):295, 2022.

33 Tadeo Saldanõ, Nahuel Escobedo, Julia Marchetti, Diego Javier Zea, Juan Mac Donagh, Ana Julia Velez Rueda, Eduardo Gonik, Agustina García Melani, Julieta Novomisky Nechcoff, Martín N Salas, et al. Impact of protein conformational diversity on alphafold predictions. Bioinformatics, 38(10):2742–2748, 2022.

34 Tareq Hameduh, Michal Mokry, Andrew D Miller, Zbynek Heger, and Yazan Haddad. Solvent accessibility promotes rotamer errors during protein modeling with major side-chain prediction programs. Journal of Chemical Information and Modeling, 63(14):4405–4422, 2023.

35 Natália Teruel, Vinicius Magalhaẽs Borges, and Rafael Najmanovich. Surfaces: a software to quantify and visualize interactions within and between proteins and ligands. Bioinformatics, 39(10):btad608, 2023.

36 Sarah Sirin, James R Apgar, Eric M Bennett, and Amy E Keating. Ab-bind: antibody binding mutational database for computational affinity predictions. Protein Science, 25(2):393– 409, 2016.

37 Yunyun Gao, Volker Thorn, and Andrea Thorn. Errors in structural biology are not the exception. Acta Crystallographica Section D: Structural Biology, 79(3):206–211, 2023.

38 Matthew McCallum, Nadine Czudnochowski, Laura E Rosen, Samantha K Zepeda, John E Bowen, Alexandra C Walls, Kevin Hauser, Anshu Joshi, Cameron Stewart, Josh R Dillen, et al. Structural basis of sars-cov-2 omicron immune evasion and receptor engagement. Science, 375(6583):864–868, 2022.

39 Dhiraj Mannar, James W Saville, Xing Zhu, Shanti S Srivastava, Alison M Berezuk, Katharine S Tuttle, Ana Citlali Marquez, Inna Sekirov, and Sriram Subramaniam. Sars-cov-2 omicron variant: Antibody evasion and cryo-em structure of spike protein–ace2 complex. Science, 375(6582):760–764, 2022.

40 Qin Hong, Wenyu Han, Jiawei Li, Shiqi Xu, Yifan Wang, Cong Xu, Zuyang Li, Yanxing Wang, Chao Zhang, Zhong Huang, et al. Molecular basis of receptor binding and antibody neutralization of omicron. Nature, 604(7906):546–552, 2022.

41 Andrej Šali and Tom L Blundell. Comparative protein modelling by satisfaction of spatial restraints. Journal of molecular biology, 234(3):779–815, 1993.

42 Olivier Mailhot and Rafael Najmanovich. The nrgten python package: an extensible toolkit for coarse-grained normal mode analysis of proteins, nucleic acids, small molecules and their complexes. Bioinformatics, 37(19):3369–3371, 2021.

43 Sophie M-C Gobeil, Katarzyna Janowska, Shana McDowell, Katayoun Mansouri, Robert Parks, Kartik Manne, Victoria Stalls, Megan F Kopp, Rory Henderson, Robert J Edwards, et al. D614g mutation alters sars-cov-2 spike conformation and enhances protease cleavage at the s1/s2 junction. Cell reports, 34(2), 2021.

44 Yifan Wang, Cong Xu, Yanxing Wang, Qin Hong, Chao Zhang, Zuyang Li, Shiqi Xu, Qinyu Zuo, Caixuan Liu, Zhong Huang, et al. Conformational dynamics of the beta and kappa sars-cov-2 spike proteins and their complexes with ace2 receptor revealed by cryo-em. Nature communications, 12(1):7345, 2021.

45 Yifan Wang, Caixuan Liu, Chao Zhang, Yanxing Wang, Qin Hong, Shiqi Xu, Zuyang Li, Yong Yang, Zhong Huang, and Yao Cong. Structural basis for sars-cov-2 delta variant recognition of ace2 receptor and broadly neutralizing antibodies. Nature communications, 13(1):871, 2022.

46 Matthew Crown, Natalia Teruel, Rafael Najmanovich, and Matthew Bashton. Spear: Systematic protein annotator. Bioinformatics, 38(15):3827–3829, 2022.

47 Dennis A Benson, Mark Cavanaugh, Karen Clark, Ilene Karsch-Mizrachi, James Ostell, Kim D Pruitt, and Eric W Sayers. Genbank. Nucleic acids research, 46(D1):D41–D47, 2018.

48 Allison J Greaney, Tyler N Starr, and Jesse D Bloom. An antibody-escape estimator for mutations to the sars-cov-2 receptor-binding domain. Virus evolution, 8(1):veac021, 2022.

49 Yunlong Cao, Fanchong Jian, Jing Wang, Yuanling Yu, Weiliang Song, Ayijiang Yisimayi, Jing Wang, Ran An, Xiaosu Chen, Na Zhang, et al. Imprinted sars-cov-2 humoral immunity induces convergent omicron rbd evolution. Nature, 614(7948):521–529, 2023.

50 Dami A Collier, Anna De Marco, Isabella ATM Ferreira, Bo Meng, Rawlings P Datir, Alexandra C Walls, Steven A Kemp, Jessica Bassi, Dora Pinto, Chiara Silacci-Fregni, et al. Sensitivity of sars-cov-2 b. 1.1. 7 to mrna vaccine-elicited antibodies. Nature, 593(7857):136–141, 2021.

51 Petra Mlcochova, Steven A Kemp, Mahesh Shanker Dhar, Guido Papa, Bo Meng, Isabella ATM Ferreira, Rawlings Datir, Dami A Collier, Anna Albecka, Sujeet Singh, et al. Sars-cov-2 b. 1.617. 2 delta variant replication and immune evasion. Nature, 599(7883):114–119, 2021.

52 Qianqian Li, Jiajing Wu, Jianhui Nie, Li Zhang, Huan Hao, Shuo Liu, Chenyan Zhao, Qi Zhang, Huan Liu, Lingling Nie, et al. The impact of mutations in sars-cov-2 spike on viral infectivity and antigenicity. Cell, 182(5):1284–1294, 2020.

53 Yao Fan, Xiang Li, Lei Zhang, Shu Wan, Long Zhang, and Fangfang Zhou. Sars-cov-2 omicron variant: recent progress and future perspectives. Signal transduction and targeted therapy, 7(1):1–11, 2022.

54 Brian J Willett, Joe Grove, Oscar A MacLean, Craig Wilkie, Giuditta De Lorenzo, Wilhelm Furnon, Diego Cantoni, Sam Scott, Nicola Logan, Shirin Ashraf, et al. Sars-cov-2 omicron is an immune escape variant with an altered cell entry pathway. Nature microbiology, 7(8):1161–1179, 2022.

55 Delphine Planas, Nell Saunders, Piet Maes, Florence Guivel-Benhassine, Cyril Planchais, Julian Buchrieser, William-Henry Bolland, Françoise Porrot, Isabelle Staropoli, Frederic Lemoine, et al. Considerable escape of sars-cov-2 omicron to antibody neutralization. Nature, 602(7898):671–675, 2022.

56 Ida Paciello, Giuseppe Maccari, Giulio Pierleoni, Federica Perrone, Giulia Realini, Marco Troisi, Gabriele Anichini, Maria Grazia Cusi, Rino Rappuoli, and Emanuele Andreano. Sars-cov-2 jn. 1 variant evasion of ighv3-53/3-66 b cell germlines. Science Immunology, 9(98):eadp9279, 2024.

57 Chiranjib Chakraborty, Manojit Bhattacharya, Hitesh Chopra, Prosun Bhattacharya, Md Aminul Islam, and Kuldeep Dhama. Recently emerged omicron subvariant bf. 7 and its r346t mutation in the rbd region reveal increased transmissibility and higher resistance to neutralization antibodies: need to understand more under the current scenario of rising cases in china and fears of driving a new wave of the covid-19 pandemic. International Journal of Surgery, 109(4):1037–1040, 2023.

58 Ramona Groenheit, Ilias Galanis, Klara Sondén, Maike Sperk, Elin Movert, Philip Bacchus, Tatiana Efimova, Lina Petersson, Marie Rapp, Viktoria Sahlén, et al. Rapid emergence of omicron sublineages expressing spike protein r346t. The Lancet Regional Health–Europe, 24, 2023.

59 Irene M Francino-Urdaniz, Paul J Steiner, Monica B Kirby, Fangzhu Zhao, Cyrus M Haas, Shawn Barman, Emily R Rhodes, Alison C Leonard, Linghang Peng, Kayla G Sprenger, et al. One-shot identification of sars-cov-2 s rbd escape mutants using yeast screening. Cell Reports, 36(9), 2021.

60 Li Zhang, Zhimin Cui, Qianqian Li, Bo Wang, Yuanling Yu, Jiajing Wu, Jianhui Nie, Ruxia Ding, Haixin Wang, Yue Zhang, et al. Ten emerging sars-cov-2 spike variants exhibit variable infectivity, animal tropism, and antibody neutralization. Communications biology, 4(1):1196, 2021.

61 Panke Qu, John P Evans, Julia N Faraone, Yi-Min Zheng, Claire Carlin, Mirela Anghelina, Patrick Stevens, Soledad Fernandez, Daniel Jones, Gerard Lozanski, et al. Enhanced neutralization resistance of sars-cov-2 omicron subvariants bq. 1, bq. 1.1, ba. 4.6, bf. 7, and ba. 2.75. 2. Cell host & microbe, 31(1):9–17, 2023.

62 Elisabetta Cameroni, John E Bowen, Laura E Rosen, Christian Saliba, Samantha K Zepeda, Katja Culap, Dora Pinto, Laura A VanBlargan, Anna De Marco, Julia di Iulio, et al. Broadly neutralizing antibodies overcome sars-cov-2 omicron antigenic shift. Nature, 602(7898):664–670, 2022.

63 Pengcheng Han, Linjie Li, Sheng Liu, Qisheng Wang, Di Zhang, Zepeng Xu, Pu Han, Xiaomei Li, Qi Peng, Chao Su, et al. Receptor binding and complex structures of human ace2 to spike rbd from omicron and delta sars-cov-2. Cell, 185(4):630–640, 2022.

64 Michael I Barton, Stuart A MacGowan, Mikhail A Kutuzov, Omer Dushek, Geoffrey John Barton, and P Anton Van Der Merwe. Effects of common mutations in the sars-cov-2 spike rbd and its ligand, the human ace2 receptor on binding affinity and kinetics. Elife, 10:e70658, 2021.

65 Tyler N Starr, Allison J Greaney, Sarah K Hilton, Daniel Ellis, Katharine HD Crawford, Adam S Dingens, Mary Jane Navarro, John E Bowen, M Alejandra Tortorici, Alexandra C Walls, et al. Deep mutational scanning of sars-cov-2 receptor binding domain reveals constraints on folding and ace2 binding. cell, 182(5):1295–1310, 2020.

66 Fang Tian, Bei Tong, Liang Sun, Shengchao Shi, Bin Zheng, Zibin Wang, Xianchi Dong, and Peng Zheng. N501y mutation of spike protein in sars-cov-2 strengthens its binding to receptor ace2. elife, 10:e69091, 2021.

67 Charlie Laffeber, Kelly de Koning, Roland Kanaar, and Joyce HG Lebbink. Experimental evidence for enhanced receptor binding by rapidly spreading sars-cov-2 variants. Journal of molecular biology, 433(15):167058, 2021.

68 Qibin Geng, Ke Shi, Gang Ye, Wei Zhang, Hideki Aihara, and Fang Li. Structural basis for human receptor recognition by sars-cov-2 omicron variant ba. 1. Journal of Virology, 96(8):e00249–22, 2022.

69 Alief Moulana, Thomas Dupic, Angela M Phillips, Jeffrey Chang, Serafina Nieves, Anne A Roffler, Allison J Greaney, Tyler N Starr, Jesse D Bloom, and Michael M Desai. Compensatory epistasis maintains ace2 affinity in sars-cov-2 omicron ba. 1. Nature Communications, 13(1):7011, 2022.

70 Maren Schubert, Federico Bertoglio, Stephan Steinke, Philip Alexander Heine, Mario Alberto Ynga-Durand, Henrike Maass, Josè Camilla Sammartino, Irene Cassaniti, Fanglei Zuo, Likun Du, et al. Human serum from sars-cov-2-vaccinated and covid-19 patients shows reduced binding to the rbd of sars-cov-2 omicron variant. BMC medicine, 20(1):102, 2022.

71 Wei Zhang, Ke Shi, Qibin Geng, Morgan Herbst, Michael Wang, Linfen Huang, Fan Bu, Bin Liu, Hideki Aihara, and Fang Li. Structural evolution of sars-cov-2 omicron in human receptor recognition. Journal of virology, 97(8):e00822–23, 2023.

72 Yunlong Cao, Ayijiang Yisimayi, Fanchong Jian, Weiliang Song, Tianhe Xiao, Lei Wang, Shuo Du, Jing Wang, Qianqian Li, Xiaosu Chen, et al. Ba. 2.12. 1, ba. 4 and ba. 5 escape antibodies elicited by omicron infection. Nature, 608(7923):593–602, 2022.

73 Izumi Kimura, Daichi Yamasoba, Hesham Nasser, Jiri Zahradnik, Yusuke Kosugi, Jiaqi Wu, Kayoko Nagata, Keiya Uriu, Yuri L Tanaka, Jumpei Ito, et al. The sars-cov-2 spike s375f mutation characterizes the omicron ba. 1 variant. Iscience, 25(12), 2022.

74 Seonghan Kim, Yi Liu, Matthew Ziarnik, Sangjae Seo, Yiwei Cao, X Frank Zhang, and Wonpil Im. Binding of human ace2 and rbd of omicron enhanced by unique interaction patterns among sars-cov-2 variants of concern. Journal of computational chemistry, 44(4):594– 601, 2023.

75 Sophie M-C Gobeil, Katarzyna Janowska, Shana McDowell, Katayoun Mansouri, Robert Parks, Victoria Stalls, Megan F Kopp, Kartik Manne, Dapeng Li, Kevin Wiehe, et al. Effect of natural mutations of sars-cov-2 on spike structure, conformation, and antigenicity. Science, 373(6555):eabi6226, 2021.

76 Sophie M-C Gobeil, Rory Henderson, Victoria Stalls, Katarzyna Janowska, Xiao Huang, Aaron May, Micah Speakman, Esther Beaudoin, Kartik Manne, Dapeng Li, et al. Structural diversity of the sars-cov-2 omicron spike. Molecular cell, 82(11):2050–2068, 2022.

77 Natalia Teruel, Matthew Crown, Matthew Bashton, and Rafael Najmanovich. Computational analysis of the effect of sars-cov-2 variant omicron spike protein mutations on dynamics, ace2 binding and propensity for immune escape. *bioRxiv*, pages 2021–12, 2021.

78 Zhennan Zhao, Jingya Zhou, Mingxiong Tian, Min Huang, Sheng Liu, Yufeng Xie, Pu Han, Chongzhi Bai, Pengcheng Han, Anqi Zheng, et al. Omicron sars-cov-2 mutations stabilize spike up-rbd conformation and lead to a non-rbm-binding monoclonal antibody escape. Nature communications, 13(1):4958, 2022.

79 Chiranjib Chakraborty, Manojit Bhattacharya, Hitesh Chopra, Md Aminul Islam, Gutulla Saikumar, and Kuldeep Dhama. The SARS-CoV-2 Omicron recombinant subvariants XBB, XBB. 1, and XBB. 1.5 are expanding rapidly with unique mutations, antibody evasion, and immune escape properties–an alarming global threat of a surge in COVID-19 cases again? International Journal of Surgery, 109(4):1041–1043, 2023.

80 Pei Li, Julia N Faraone, Cheng Chih Hsu, Michelle Chamblee, Yi-Min Zheng, Claire Carlin, Joseph S Bednash, Jeffrey C Horowitz, Rama K Mallampalli, Linda J Saif, et al. Characteristics of jn. 1-derived sars-cov-2 subvariants slip, flirt, and kp. 2 in neutralization escape, infectivity and membrane fusion. BioRxiv, 2024.

81 Oliver C Grant, David Montgomery, Keigo Ito, and Robert J Woods. Analysis of the sarscov-2 spike protein glycan shield reveals implications for immune recognition. Scientific reports, 10(1):14991, 2020.

82 Wilton B Williams, R Ryan Meyerhoff, RJ Edwards, Hui Li, Kartik Manne, Nathan I Nicely, Rory Henderson, Ye Zhou, Katarzyna Janowska, Katayoun Mansouri, et al. Fab-dimerized glycan-reactive antibodies are a structural category of natural antibodies. Cell, 184(11):2955–2972, 2021.

83 Priyamvada Acharya, Wilton Williams, Rory Henderson, Katarzyna Janowska, Kartik Manne, Robert Parks, Margaret Deyton, Jordan Sprenz, Victoria Stalls, Megan Kopp, et al. A glycan cluster on the sars-cov-2 spike ectodomain is recognized by fab-dimerized glycan-reactive antibodies. BioRxiv, pages 2020–06, 2020.

84 Kien Nguyen, Srirupa Chakraborty, Rachael A Mansbach, Pedro D Manrique, Bette Korber, and Sandrasegaram Gnanakaran. Exploring the role of glycans in the interaction of sars-cov2 rbd and human receptor ace2. Biophysical Journal, 120(3):15a, 2021.

85 Joel D Allen, Yasunori Watanabe, Himanshi Chawla, Maddy L Newby, and Max Crispin. Subtle influence of ace2 glycan processing on sars-cov-2 recognition. Journal of molecular biology, 433(4):166762, 2021.

86 Yanqiu Gong, Suideng Qin, Lunzhi Dai, and Zhixin Tian. The glycosylation in sars-cov-2 and its receptor ace2. Signal Transduction and Targeted Therapy, 6(1):396, 2021.

87 Terra Sztain, Surl-Hee Ahn, Anthony T Bogetti, Lorenzo Casalino, Jory A Goldsmith, Evan Seitz, Ryan S McCool, Fiona L Kearns, Francisco Acosta-Reyes, Suvrajit Maji, et al. A glycan gate controls opening of the sars-cov-2 spike protein. Nature chemistry, 13(10):963– 968, 2021.

88 Yui Tik Pang, Atanu Acharya, Diane L Lynch, Anna Pavlova, and James C Gumbart. Sars-cov-2 spike opening dynamics and energetics reveal the individual roles of glycans and their collective impact. Communications Biology, 5(1):1170, 2022.

89 Aoife M Harbison, Carl A Fogarty, Toan K Phung, Akash Satheesan, Benjamin L Schulz, and Elisa Fadda. Fine-tuning the spike: role of the nature and topology of the glycan shield in the structure and dynamics of the sars-cov-2 s. Chemical Science, 13(2):386–395, 2022.

90 Shuyuan Zhang, Qingtai Liang, Xinheng He, Chongchong Zhao, Wenlin Ren, Ziqing Yang, Ziyi Wang, Qiang Ding, Haiteng Deng, Tong Wang, et al. Loss of spike n370 glycosylation as an important evolutionary event for the enhanced infectivity of sars-cov-2. Cell research, 32(3):315–318, 2022.

91 Chaoran Chen, Sarah Nadeau, Michael Yared, Philippe Voinov, Ning Xie, Cornelius Roemer, and Tanja Stadler. Cov-spectrum: analysis of globally shared sars-cov-2 data to identify and characterize new variants. Bioinformatics, 38(6):1735–1737, 2022.

92 Sören von Bülow, Mateusz Sikora, Florian EC Blanc, Roberto Covino, and Gerhard Hummer. Antibody accessibility determines location of spike surface mutations in sars-cov-2 variants. PLOS Computational Biology, 19(1):e1010822, 2023.

93 Luping Zheng, Ke Wang, Minghai Chen, Fujun Qin, Chuang Yan, and Xian-En Zhang. Characterization and function of glycans on the spike proteins of sars-cov-2 variants of concern. Microbiology Spectrum, 10(6):e03120–22, 2022.

94 Sabyasachi Baboo, Jolene K Diedrich, Jonathan L Torres, Jeffrey Copps, Bhavya Singh, Patrick T Garrett, Andrew B Ward, James C Paulson, and John R Yates III. Evolving spike-protein n-glycosylation in sars-cov-2 variants. Biorxiv, 2023.

95 Yaning Li, Chang Ren, Yaping Shen, Yuanyuan Zhang, Jin Chen, Jiangnan Zheng, Ruijun Tian, Liwei Cao, and Renhong Yan. Cryo-em structures of sars-cov-2 ba. 2-derived subvariants spike in complex with ace2 receptor. Cell Discovery, 9(1):108, 2023.

96 Asif Shajahan, Lauren E Pepi, Bhoj Kumar, Nathan B Murray, and Parastoo Azadi. Site specific n-and o-glycosylation mapping of the spike proteins of sars-cov-2 variants of concern. Scientific Reports, 13(1):10053, 2023.

97 Pan Liu, Can Yue, Bo Meng, Tianhe Xiao, Sijie Yang, Shuo Liu, Fanchong Jian, Qianhui Zhu, Yuanling Yu, Yanyan Ren, et al. Spike n354 glycosylation augments sars-cov-2 fitness for human adaptation through structural plasticity. National Science Review, page nwae206, 2024.

98 Justin Bryan Goh and Say Kong Ng. Impact of host cell line choice on glycan profile. Critical reviews in biotechnology, 38(6):851–867, 2018.

99 Weiwei Li, Zepeng Xu, Tianhui Niu, Yufeng Xie, Zhennan Zhao, Dedong Li, Qingwen He, Wenqiao Sun, Kaiyuan Shi, Wenjing Guo, et al. Key mechanistic features of the trade-off between antibody escape and host cell binding in the sars-cov-2 omicron variant spike proteins. The EMBO Journal, 43(8):1484–1498, 2024.

100 Drew Weissman, Mohamad-Gabriel Alameh, Thushan de Silva, Paul Collini, Hailey Hornsby, Rebecca Brown, Celia C LaBranche, Robert J Edwards, Laura Sutherland, Sampa Santra, et al. D614g spike mutation increases sars cov-2 susceptibility to neutralization. Cell host & microbe, 29(1):23–31, 2021.

