## Supplementary Tables and Images for "Comprehensive Analysis of SARS-CoV-2 Spike Evolution: Epitope Classification and Immune Escape Prediction"

### Supplementary Material

**Table S1 Conservation of the most frequently glycosylated positions with described functional aspects.** Data according to CoV-Spectrum as of August 15, 2024.

| Position | Glycosylation in the structural dataset | Functional observation | Number of mutated sequences | Overall proportion |
|---|---|---|---|---|
| 343 | 1,794 | Antibody recognition and Inter-chain interactions | 13,588 | 0.08% |
| 709 | 1,624 | Antibody recognition and Inter-chain interactions | 11,048 | 0.07% |
| 801 | 1,563 | Antibody recognition | 20,774 | 0.13% |
| 717 | 1,555 | Antibody recognition | 11,362 | 0.07% |
| 282 | 1,535 | Antibody recognition and Inter-chain interactions | 17,716 | 0.11% |
| 1074 | 1,528 | Inter-chain interactions | 214,511 | 1.3% |
| 61 | 1,499 | Antibody recognition | 17,927 | 0.11% |
| 616 | 1,486 | Inter-chain interactions | 7,488 | 0.05% |
| 331 | 1,474 | Antibody recognition | 14,687 | 0.09% |
| 234 | 1,255 | Antibody recognition and Inter-chain interactions | 16,857 | 0.1% |
| 165 | 1,172 | Antibody recognition and Inter-chain interactions | 17,298 | 0.11% |
| 122 | 1,121 | Antibody recognition | 17,123 | 0.1% |
| 603 | 1,099 | Antibody recognition | 7,397 | 0.04% |
| 17 | 393 | Antibody recognition | 43,164 | 0.26% |
| 149 | 229 | Antibody recognition | 17,045 | 0.1% |

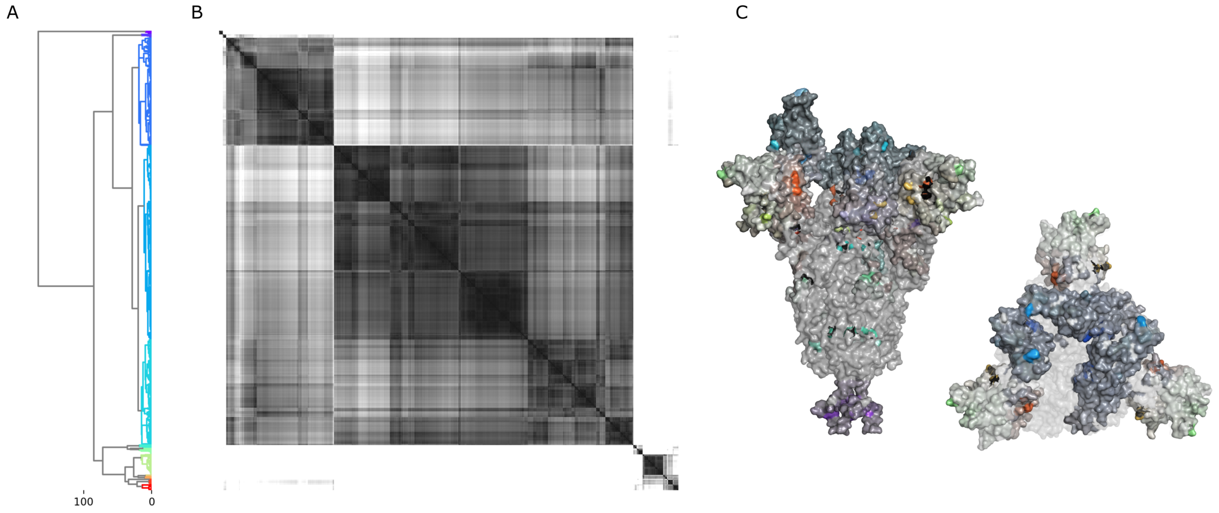


**Fig S1 Initial epitope clustering.** (A) Dendrogram of the hierarchical clustering of 2032 vectors of interaction of antibodies according to their net interaction per Spike residue. (B) Euclidean distance matrix of the average points of interaction of each antibody, cut by dashed lines according to their respective dendrogram clusters, defined by a distance threshold of 19Å. (C) Structural representation of 14 separated branches according to the colors of the dendrogram.

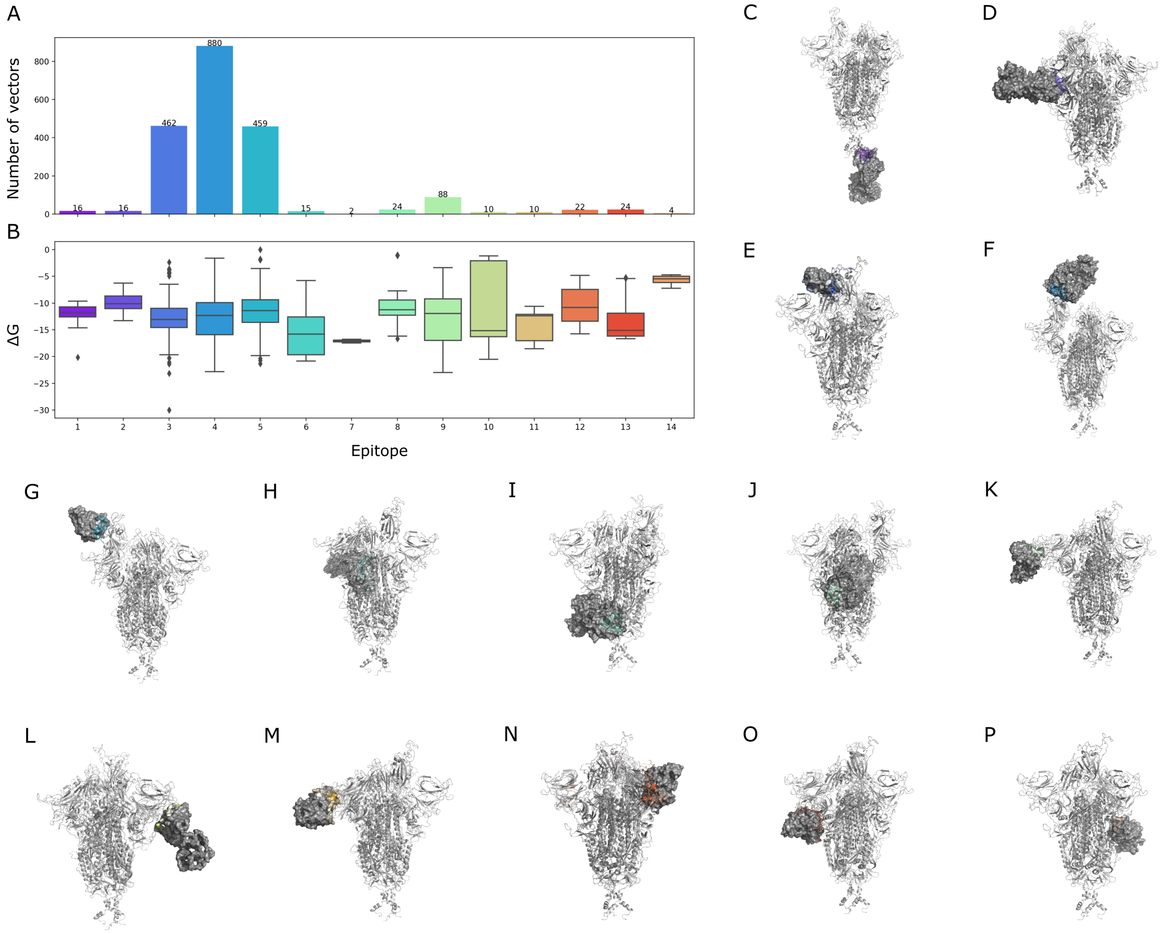


**Fig S2 Detailed characterization of the 14 epitopes.** (A) Number of interaction vectors associated with each epitope. (B) Calculated binding affinity ($\Delta G$, kcal/mol) for each epitope. Representatives of average-binding antibodies targeting each epitope, following the same epitope color coding, are (C) chains A and B from PDB 7NAB for epitope 1, (D) chains I and J from PDB 7WD9 for epitope 2, (E) chain B from PDB 7FBJ for epitope 3, (F) chains H and L from PDB 7KXK for epitope 4, (G) chains A and B from PDB 7V27 for epitope 5, (H) chains A and B from PDB 8GOM for epitope 6, (I) chains A and B from PDB 7M8U for epitope 7, (J) chains E and F from PDB 8D6Z for epitope 8, (K) chains H and L from PDB 7RQ6 for epitope 9, (L) chains O and P from PDB 7DZY for epitope 10, (M) chains O and P from PDB 7UAP for epitope 11, (N) chains E and F from PDB 7SOB for epitope 12, (O) chains D and E from PDB 7UAR for epitope 13, and (P) chain D from PDB 7JJC for epitope 14.

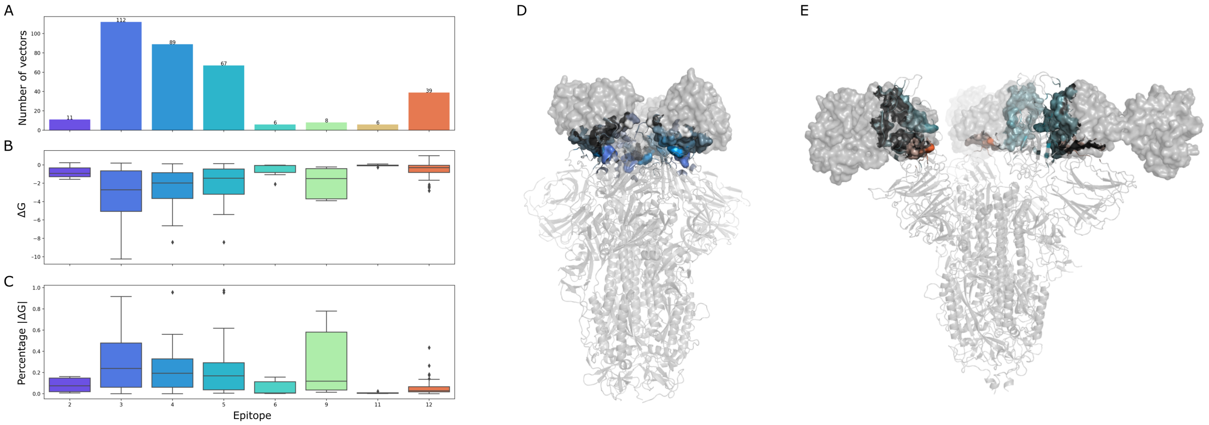


**Fig S3 Detailed characterization of secondary interactions.** (A) Number of secondary interaction vectors associated with each epitope. (B) Calculated binding affinity ($\Delta G$, kcal/mol) for each epitope. (C) Percentage of binding affinity of that of the main interaction. (D) Interaction interface of the antibody S2M11, mainly interacting with epitope 4 and also interacting with the epitope 3 of the neighbouring chain (PDB 8DLW).(E) Interaction interface of the antibody R1-32, mainly interacting with epitope 5 and also interacting with the epitope 12 of the neighbouring chain (PDB 7YE9).

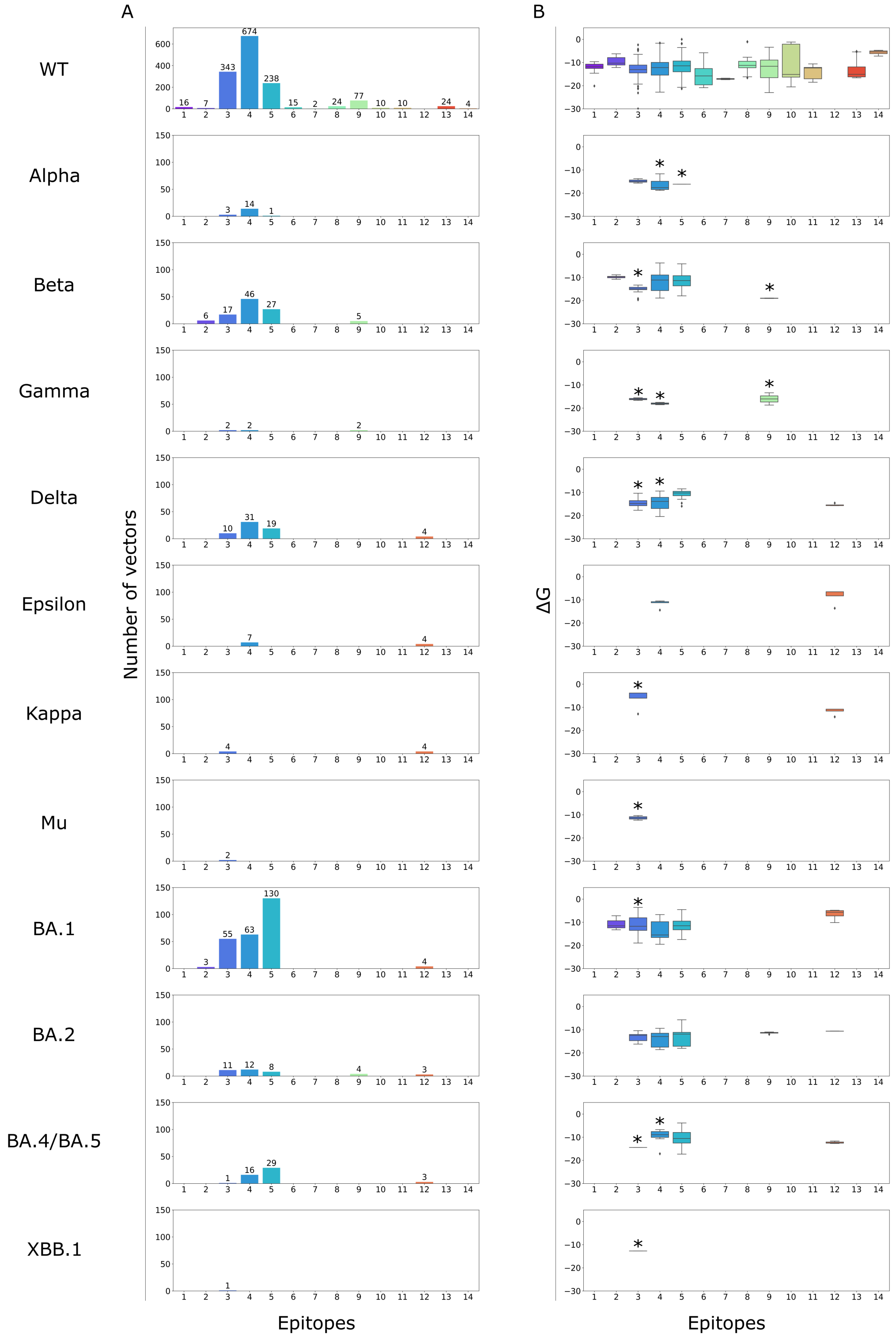

**Fig S4 Epitope analysis across variants of concern compared to the *wild-type*.** (A) Number of interaction vectors associated with each epitope for each variant. (B) Calculated binding affinity ($\Delta G$, kcal/mol) for each epitope group.

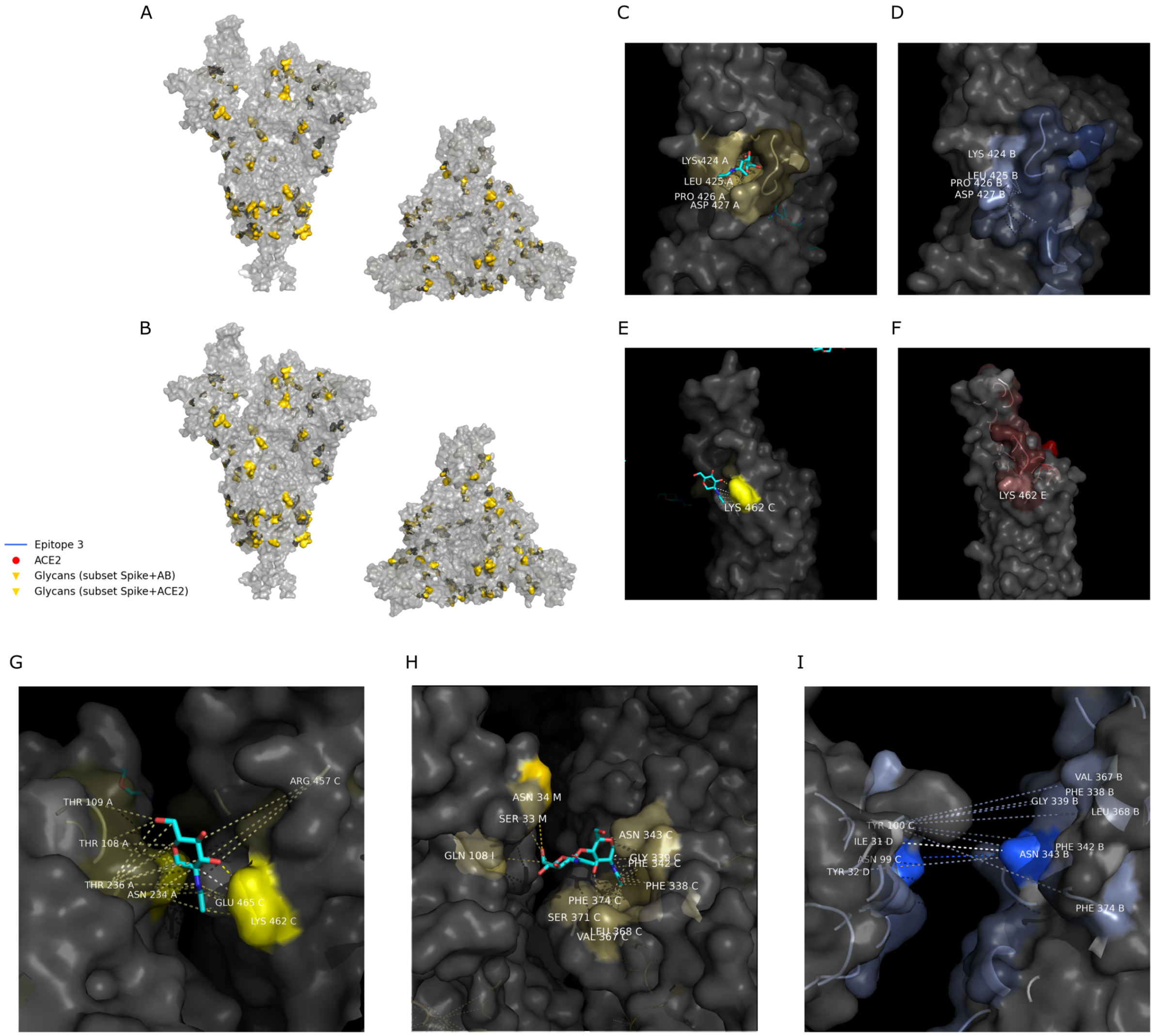


**Fig S5 Detailed structural representation of the glycan coated residues.** (A) Structural representation of the glycan coated residues based on the evaluation of Spike structures in complex with antibodies (gold), (B) or Spike structures in complex with ACE2 (yellow); these residues are either directly glycosylated, or in neighboring areas and with existing interactions with glycans. (C) We can see glycans interacting with residues such as LYS424, LEU425, PRO426 and ASP427 (PDB 7YQX), (D) which in other structures are also seen as part of the antibody binding surface for epitope 3 (PDB 8CYC). (E) Among the structures that are in complex with ACE2, we see strong glycan interactions with LYS462 (PDB 8DM9), (F) a residue that is also part of the interacting interface with ACE2 (PDB 7F5R). (G) The context of the LYS462 interacting glycan is a covalent interaction to the ASN234 of the neighboring Spike chain (PDB 8DM9). (H) We can find examples of co-interactions for complexes with antibodies, such as a glycan linked to ASN343 and showing favorable interactions with heavy and light antibody chains (PDB 7CZX), (I) while same residues PHE338, GLY339, PHE342, ASN343, VAL367, LEU368 and PHE374, from epitope 3, can be found strongly interacting with antibodies without the presence of glycans (PDB 8DW9) (exploded interaction for visual purposes).

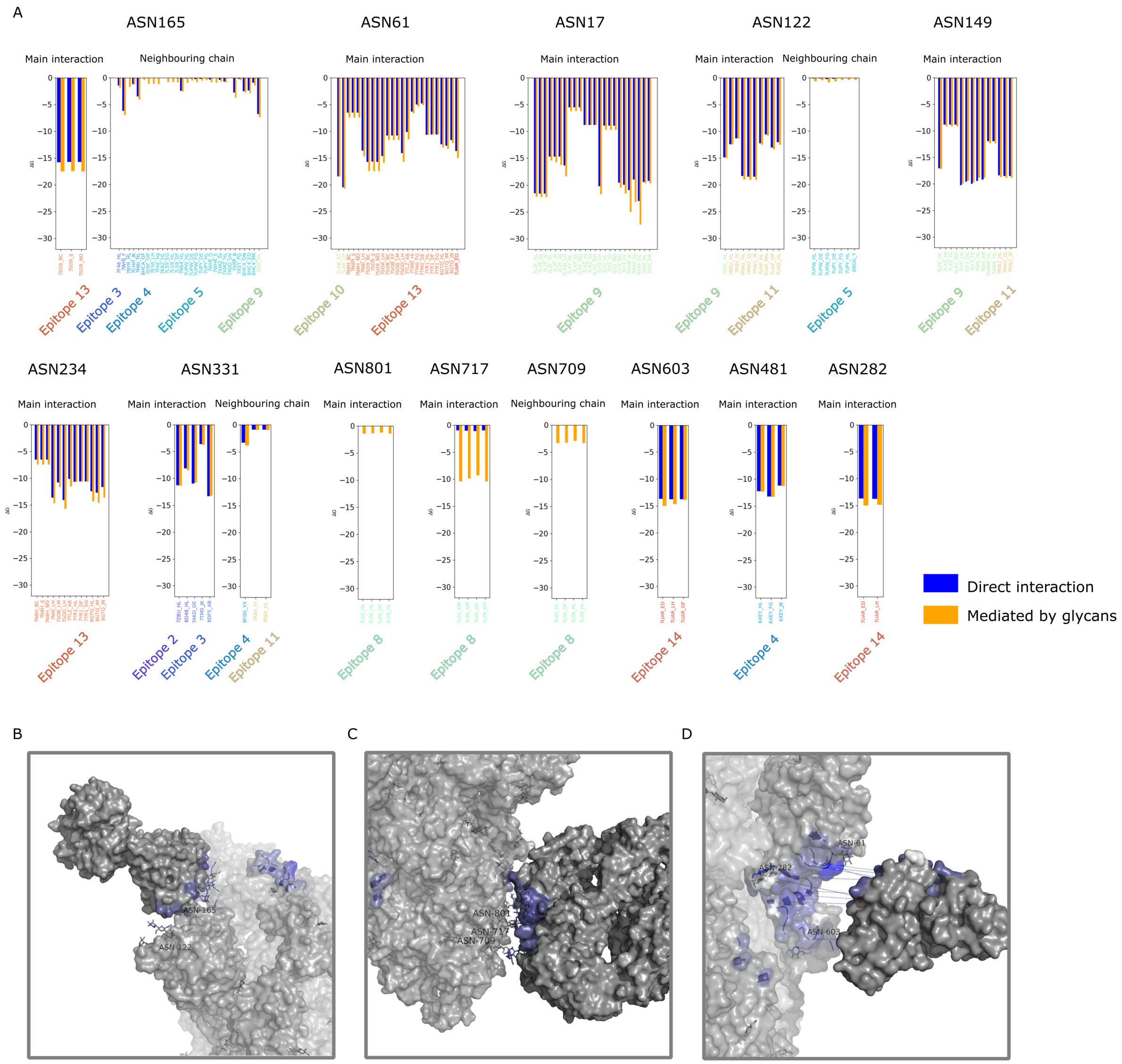


**Fig S6 Compared binding affinity for glycan-mediated antibody interactions.** (A) Values of binding affinity ($\Delta G$, kcal/mol) comparing direct interaction (blue) to interaction mediated by glycans (orange) linked to ASN165, ASN61, ASN17, ASN122, ASN149, ASN234, ASN331, ASN801, ASN717, ASN709, ASN603, ASN481 and ASN282, both for co-interactions in which the glycan mediates binding of antibody chains to its main interacting Spike chain, or to a neighboring Spike chain. (B) Example of a glycan-mediated interaction in which ASN122 and ASN165-linked glycans enhance the interaction with the antibody SP1-77, targeting the neighboring epitope 5 (PDB 7UPY). (C) Representation of the glycan-dependent interaction of antibody 2G12, mediated by glycans linked to ASN709, ASN717 and ASN801 (PDB 7L06). (D) Antibody C1717, targeting epitope 13 and with increased interaction mediated by glycans linked to residues ASN61, ASN282 and ASN603 (PDB 7UAR) (exploded interaction for visual purposes).

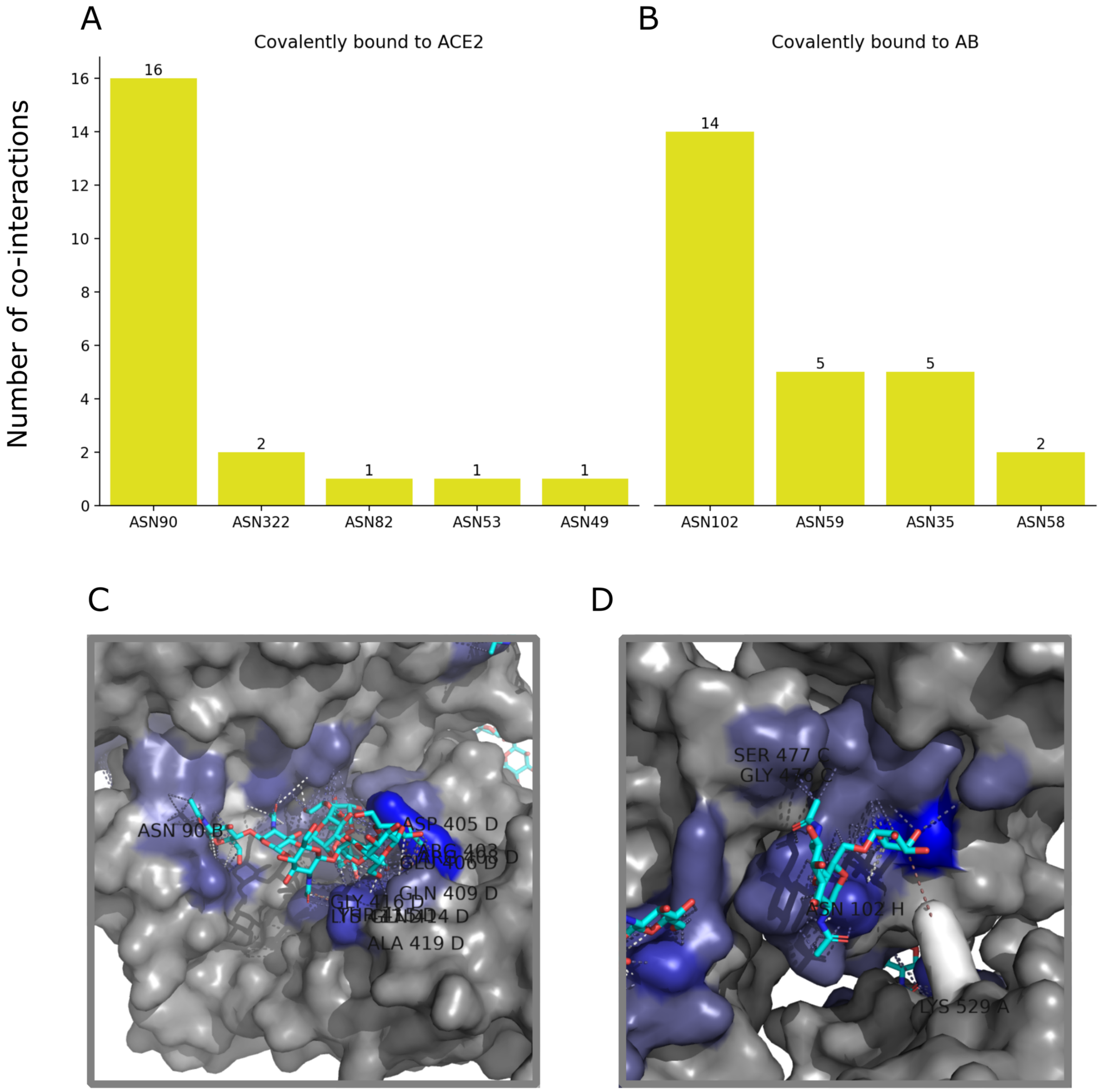


**Fig S7 Dataset representation of glycans linked to proteins other than the Spike protein.** (A) Number of co-interacting glycans based on the Spike residue they are covalently linked to in ACE2 chains and (B) antibody chains. (C) Example of glycan linked to residue ASN90 from ACE2 chain B and co-interacting with many residues from Spike chain D (PDB 7SN0). (D) Example structure of a glycan linked to residue ASN102 from antibody COVOX-253H55L, labeled as chain H, and co-interacting with chains A and C from the Spike protein (PDB 7NDA).
